# Diverse activity in prefrontal projections promotes temporal control of action

**DOI:** 10.1101/2025.05.12.653607

**Authors:** Xin Ding, Matthew A. Weber, Trevor C. Butler, Alexandra S. Bova, Stephanie Guerrero, Christopher Hunter, Rachel C. Cole, Hannah R. Stutt, Madison McMurrin, Mackenzie M. Spicer, Mackenzie M. Conlon, Shane A. Heiney, Jon M. Resch, Nandakumar S. Narayanan

## Abstract

Prefrontal neurons can have diverse activity during cognitive functions like working memory, attention, and timing; however, the importance of this heterogeneity is unclear. Our goal was to better understand the diversity of prefrontal activity through connectivity. We harnessed circuit-specific tools to capture activity within prefrontal projections during interval timing, an elementary cognitive process that requires working memory for temporal rules and attention to the passage of time to estimate a temporal interval of several seconds. We used human electroencephalography and single neuronal recordings in mice to capture prefrontal activity during interval timing, with major patterns characterized by time-dependent ramping (monotonic changes) over a temporal interval. We then leveraged retrograde viruses to interrogate prefrontal cortex (PFC) projections to the mediodorsal thalamus (PFC-MD) and to the dorsomedial striatum (PFC-DMS). We report three novel results. First, circuit-specific calcium fiber photometry revealed that PFC-MD and PFC-DMS activity encoded distinct temporal signals, with PFC-MD projections ramping down and PFC-DMS ramping up to interval timing response times. Second, circuit-specific inactivation revealed that PFC-DMS inactivation disrupted animals’ internal estimates of time. Third, circuit-specific single-nucleus RNA sequencing of prefrontal projections revealed distinct transcriptomic profiles between PFC-MD and PFC-DMS projections, with enriched genes for cortical layers and neuromodulators, and specific genes such as *Cux2, Camk2n1, Htr4,* and *Foxp2*. These data suggest differences in gene expression and connectivity give rise to the diversity of prefrontal activity during interval timing. These findings advance our fundamental understanding of prefrontal function and dysfunction in human disease.

## Introduction

Diversity enhances the function of complex systems. The prefrontal cortex (PFC) controls higher-order executive functions such as working memory, attention, and timing^1–3^ and has an incredible diversity of activity patterns. However, the significance of this different activity patterns is unclear. Understanding PFC diversity is important, particularly in the context of emerging molecular insight into prefrontal networks^4^ and for the development of treatments for diseases that affect prefrontal function, such as Parkinson’s disease and schizophrenia^5,6^.

Underlying the complexity of PFC is that prefrontal neurons are connected with a broad range of brainstem, subcortical, and cortical regions^7–9^. We characterized PFC diversity by studying two major projections to the thalamus and striatum. Projections to these areas are involved in working memory, reward learning, and inhibitory control^10–12^; yet it is unknown what information is encoded by the activity of these prefrontal projections. Here, we studied prefrontal projections during an elementary executive function, interval timing, which requires participants to estimate a temporal interval of several seconds by making a motor response. Interval timing is ideal for investigating the information encoded by prefrontal projections because 1) timing requires working memory for temporal rules and attention to the passage of time^13^, 2) timing engages prefrontal circuits^3,14–18^, 3) timing is translationally relevant for human diseases^19–22^, and 4) prefrontal neurons encode time through “ramping” activity, i.e., monotonic increases or decreases in neuronal firing rates across temporal intervals^17,18,23,24^. Ramping is ubiquitous in frontal circuits and can be captured by drift-diffusion models that integrate the accumulation of temporal evidence^23,25–27^. PFC ramping can be diverse, with the activity of single prefrontal neurons ramping up or down. Despite these data, it is unclear which prefrontal neuronal populations ramp, whether this ramping activity increases or decreases, and how ramping activity maps on to prefrontal connectivity.

We interrogated circuit specific projections in prefrontal neurons that project to the mediodorsal thalamus (PFC-MD) and to the dorsomedial striatum (PFC-DMS) during interval timing. Strikingly, we found that PFC-MD and PFC-DMS projections had distinct patterns of activity and opposite ramping directions, with PFC-MD projections ramping down and PFC-DMS projections ramping up to interval timing response times. Pathway-specific terminal optogenetic inhibition found that both types of projections influenced temporal variability, but only PFC-DMS projections controlled internal estimates of time. In addition, projection-specific single-nucleus RNA sequencing (snRNA-seq) revealed that PFC-MD and PFC-DMS projections had distinct transcriptional signatures associated with cortical layers and neuromodulators, and genes such as *Cux2, Camk2n1, Htr4,* and *Foxp2*. Together, these data underscore the importance of projection-specific organization in shaping diverse functional outputs from the prefrontal cortex, suggesting that connectivity may be a key contributor to functional heterogeneity.

## Results

### Diverse PFC activity is involved in the temporal control of action

We investigated prefrontal activity during temporal control of action using an interval timing task in which humans estimated a 2-second interval. The estimated time interval was indicated by a computer keyboard response, which involved a switch from the left-arrow key to the right-arrow key (Fig 1a). The moment when this response was made was termed the switch *response time*. In 40 human participants, response times were 2.5 ± 0.02 seconds, with a coefficient of variance (CV = standard deviation/mean) of 10.1 ± 0.05% (mean ± SEM; Fig 1b–d). Human electroencephalography (EEG) recorded from anterior frontal electrodes during the interval timing task revealed prominent time-related ramping dynamics (Fig 1e; electrode AFz), consistent with past work on EEG dynamics during interval timing^28,29^. Data-driven principal component analysis (PCA) demonstrated that the first component (PC1) explained 60% of EEG variance: activity ramped down after the trial start and ramped up until the time of the keyboard response (Fig 1f). PC1 scores indicated that this time-dependent ramping was prominent over the frontal electrodes, which broadly captured prefrontal activity (Fig 1g).

**Figure 1:**
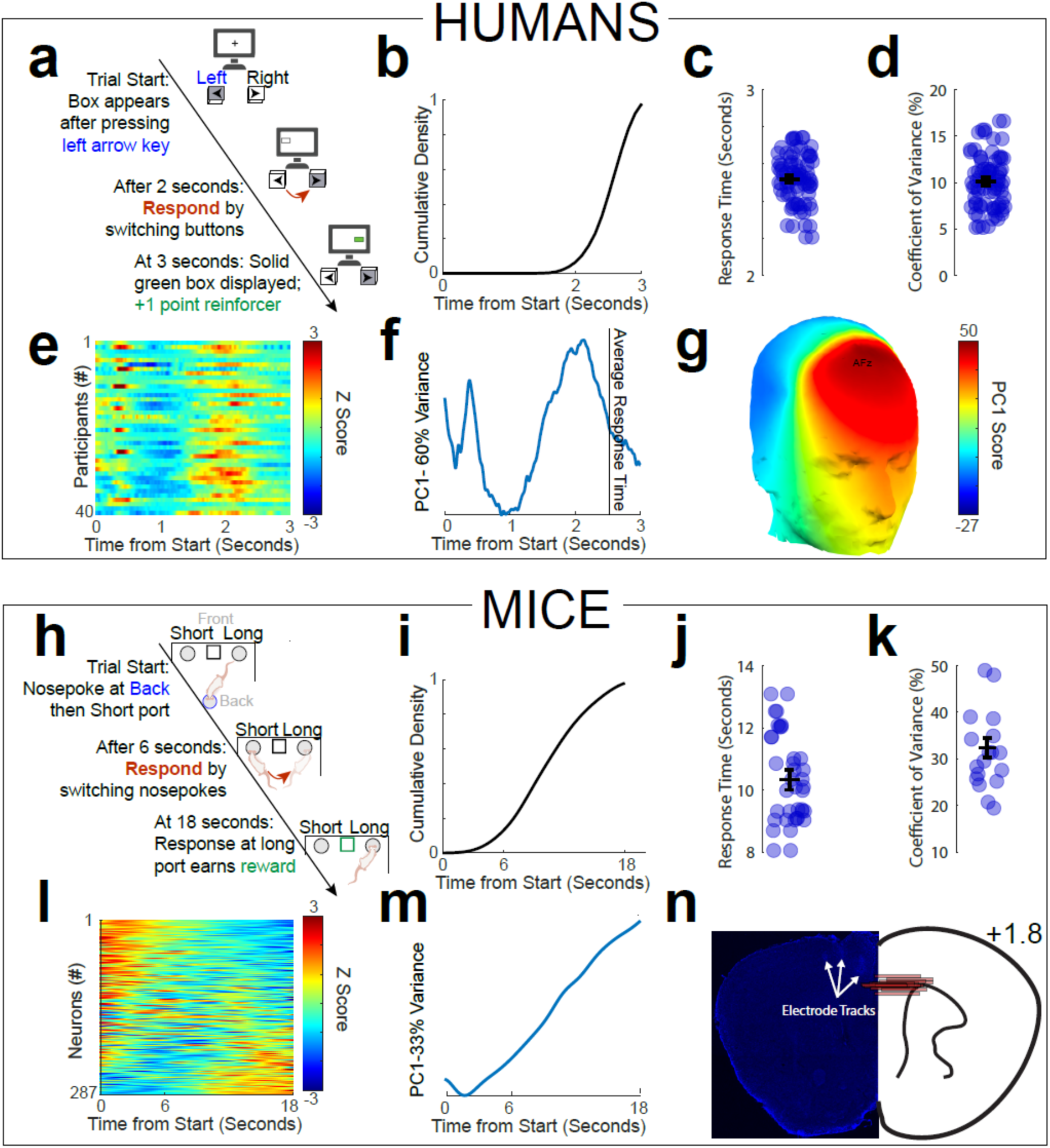
Prefrontal activity during interval timing. a) Human participants performed an interval timing task in which they were asked to switch response keys after 2 seconds but before 3 seconds to receive reinforcers. b) Cumulative distribution function of response times from 40 human participants (32 female). c) Switch response time and d) coefficient of variance (CV) from 40 human participants (32 female). e) We analyzed scalp encephalography (EEG) from the anterior frontal electrode AFz in 40 human participants during interval timing. Event-related potentials revealed highly consistent activity that ramped down after the trial start and ramped up to the time of response. f) We analyzed EEG activity using principal component analysis (PCA). The first principal component (PC1) captured ramping dynamics and explained 60% of variance from EEGs recorded during interval timing. Black line is the average switch response time. g) Scalp topography of EEG PC1 scores revealed that time-dependent ramping was focused on frontal electrodes. h) We trained 19 mice to perform a mouse interval timing task in which they switched nosepoke ports after ∼6 seconds to then receive a reward after 18 seconds. Switch response time was defined as the moment when mice depart the short nosepoke port prior to arriving at the long nosepoke port. Mice switch from the short to the long nosepoke port if they do not receive a reward at the short nosepoke port after ∼6 seconds; they receive a reward at the long nosepoke port after 18 seconds. Response times are guided by internal estimates of time and reflect temporal control of action. See Methods and Materials and Video S1 for details. i) Cumulative distribution functions of response times for 19 mice (11 female). j) Average response times and k) CV from mice. l) Single neuronal recordings from 287 prefrontal neurons in 19 mice revealed diverse time-dependent ramping activity that increased or decreased over the interval, sorted by PC1 from PCA. Neurons shown near the top of this plot appear to ramp down while neurons shown near the bottom of the plot appear to ramp up. m) Despite this diversity in ramping activity, PCA analyses of single neuronal recordings revealed prominent time-related ramping dynamics in PC1, explaining 33% variance across the prefrontal ensemble. n) Recording locations in the rodent prefrontal cortex (PFC). Estimated dorsoventral location of each recording array is shown with a red line showing the mediolateral extent of electrode arrays.

Timing is a universal feature of mammalian behavior^13^ and can be mechanistically investigated in rodent models^20,22,30,31^. We used a mouse-optimized version of an interval timing task in which mice switched responses from a “short” nosepoke port to “long” nosepoke port after ∼6 seconds to receive a food reward (Figure 1h; described in detail previously^31–37^; see Methods and Materials and Video S1). We focused on the switch *response time*, defined as the moment when a mouse exited the short nosepoke port before moving to the long nosepoke port. Since all cues are constant during the interval, mice are supposed to switch to the long nosepoke port if they are not rewarded at the short nosepoke port after ∼6 seconds. The response time is guided explicitly by the animal’s internal estimate of time and reflects temporal control of action (Fig 1h). In 19 mice, response times were 10.3 ± 0.3 seconds with a CV of 32 ± 2% (Fig 1i–k).

During interval timing trials we recorded neuronal activity from the PFC of these mice and isolated 287 prefrontal neurons across all 19 mice. We found many patterns of time-related ramping activity, with trial-by-trial linear models revealing that 36% of neurons significantly ramped down, 12% of neurons significantly ramped up, and 52% of neurons did not exhibit significant ramping activity over the 18 second interval (Fig 1l). PCA revealed that PC1 exhibited prominent ramping dynamics and explained 33% of variance across rodent prefrontal ensembles (Fig 1m–n). These data demonstrate that ramping dynamics are prominent across species in both human EEG and mouse single prefrontal neurons. We dissected the details of this activity by utilizing circuit-specific methods to isolate and interrogate PFC projections.

### Dissecting prefrontal projections

We mapped two of the largest prefrontal projections to the thalamus and the striatum^8,38^. Both of these brain regions have well-characterized roles in interval timing ^17,18,24,35,39^. Whole-brain light sheet imaging with retrograde AAV tracers (EGFP in MD and mCherry in DMS) revealed that the PFC sends spatially distinct projections to the mediodorsal thalamus and dorsomedial striatum (Video S1). PFC-MD projections were centered on deeper Layers V/VI and were dorsal and more rostral, whereas PFC-DMS projections spanned Layers II/III and V and were caudal and more ventral (Fig 2a).

**Figure 2:**
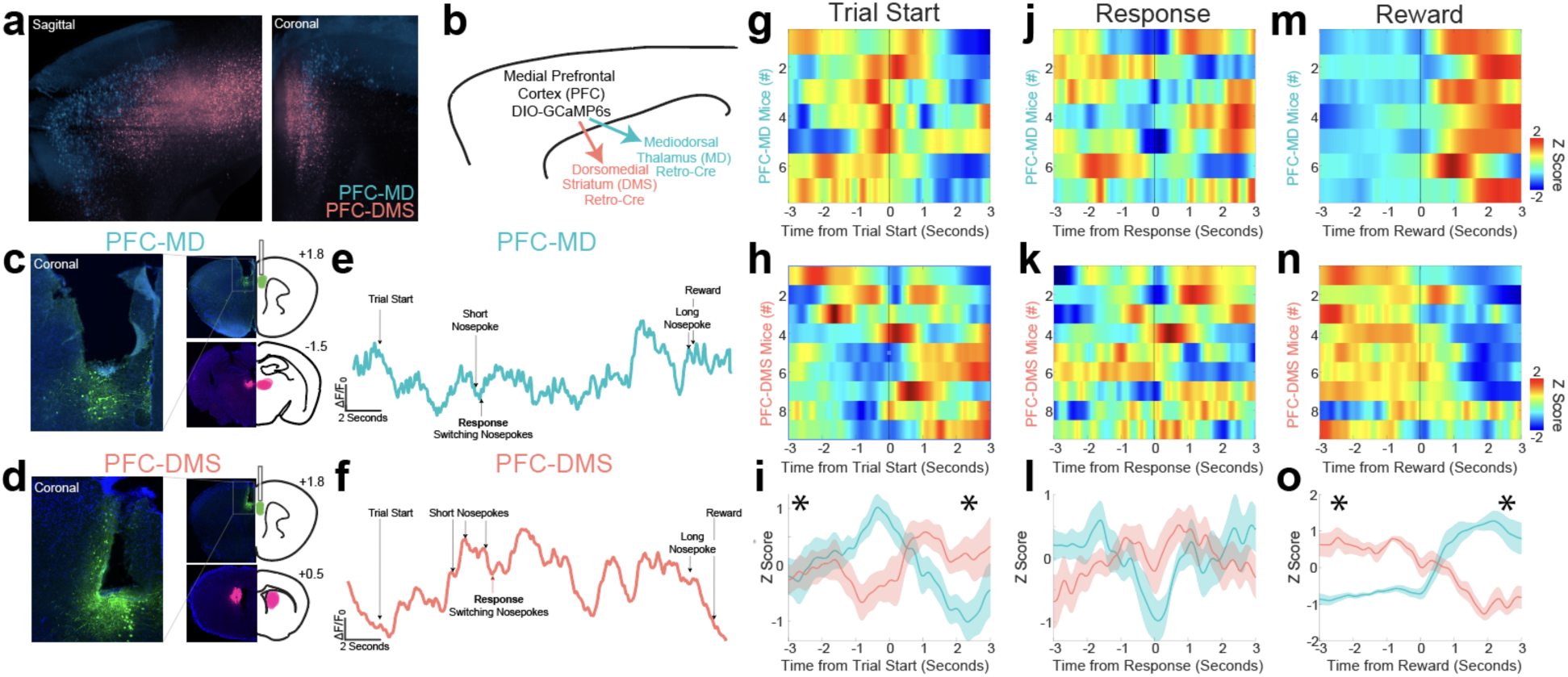
Dissecting prefrontal projections to thalamus and striatum. a) We used Retrograde AAV viruses to isolate neurons that projected from the mouse PFC to the mediodorsal thalamus (PFC-MD; aqua) and from the PFC to the dorsomedial striatum (PFC-DMS; rose). Whole-brain light sheet microscopy revealed that PFC-MD and PFC-DMS projections involved distinct prefrontal projection neurons (sagittal and coronal sections from 3D brain volume shown; Video S1). b) By injecting the Cre-dependent calcium sensor GCaMP6s into the PFC and viruses retrogradely transporting Cre-recombinase into MD or DMS, we could isolate c) PFC-MD projections or d) PFC-DMS projections; coronal section shown. Each fiber optic location with GCaMP6s virus expression is shown with a green dot; Retro-Cre viral projection fields are shown in fuchsia. We recorded activity with PFC-MD and PFC-DMS projections using GCaMP. Activity from both e) PFC-MD and f) PFC-DMS projections was prominently modulated during interval timing, as shown by ΔF/F_0_ traces from a single interval timing trial. Trial events are marked with arrows. average activity from g) PFC-MD and h) PFC-DMS animals around trial start; each row is the average z-scored activity from each animal; i) average activity across mice ± SEM. average activity from j) PFC-MD and k) PFC-DMS animals around switch response; each row is the average z-scored activity from each animal; l) average activity across mice ± SEM. average activity from m) PFC-MD and n) PFC-DMS animals around reward delivery; each row is the average z-scored activity from each animal; o) average activity across mice ± SEM. \**p* < 0.05 via a paired t-test for −3–0 seconds or 0–3 seconds. Data from 2 sessions in 7 PFC-MD mice (4 female) and 9 PFC-DMS mice (4 female).

### PFC projections have distinct patterns of activity around trial start and reward

We combined circuit-specific retrograde tools with calcium imaging using GCaMP6s fiber photometry to record activity from populations of PFC-MD or PFC-DMS projecting neurons (Fig 2b–d). Activity from PFC-MD and PFC-DMS projections was prominently modulated during interval timing (Fig 2e–f). We aligned calcium activity for PFC-MD and PFC-DMS projections around key task events such as trial start (Fig 2g–i), switch responses (Fig 2j–l), and rewards (Fig 2m–o). We included 7 mice for PFC-MD fiber photometry (Fig 2g,j,m) and 9 mice for PFC-DMS fiber photometry (Fig 2h,k,n). We found markedly distinct patterns of activity in PFC-MD and PFC-DMS projections. For instance, PFC-MD activity was greater than PFC-DMS activity before the trial start (t_(13)_ = 3.4, *p* = 0.005; −3–0 seconds before start), but less than PFC-DMS activity after the trial start (t_(13)_ = −3.4, *p* = 0.005; 0–3 seconds after start; Fig 2i). By contrast, there were few reliable differences in PFC-MD and PFC-DMS activity around the response time (t_(13)_ = 0.43, *p* = 0.68; 0–3 seconds before the switch response; t_(13)_ = −0.43, *p* = 0.67; 0–3 seconds after the switch response; Fig 2l), suggesting that motor preparatory activity was not grossly distinct between PFC-MD and PFC-DMS. Activity around nosepoke entries was not different between PFC-MD and PFC-DMS activity (Fig S1). However, PFC-MD activity was less than PFC-DMS activity around rewards, which were delivered for long nosepoke port entries after 18 seconds (t_(13)_ = −7.6, *p* = 0.5 × 10^−5^; −3–0 seconds before reward) and greater than PFC-DMS after reward delivery (t_(13)_ = 7.6, *p* = 0.5 × 10^−5^; 0–3 seconds after reward; Fig 2o). These data suggest that PFC-MD and PFC-DMS projections had distinct patterns of activity during task-related events.

### PFC-MD and PFC-DMS projections have opposite ramping activity during interval timing

Next, we examined PFC-MD and PFC-DMS projections over the temporal interval and found that they had distinct activity around the trial start and reward. As shown in Figure 1, prefrontal activity in humans and rodents exhibits time-related ramping dynamics—consistent increases or decreases in activity over a temporal interval^23,40^. We recorded activity from PFC-MD (Fig 3a) and PFC-DMS (Fig 3b) during mouse interval timing and found that PFC-MD activity decreased over the interval, whereas PFC-DMS activity increased over the interval. This difference in ramping dynamics was apparent across GCaMP activity from PFC-MD (Fig 3c) and PFC-DMS (Fig 3d) projections, and strikingly, in average activity from PFC-MD (Fig 3e) and PFC-DMS (Fig 3f) projections.

**Figure 3:**
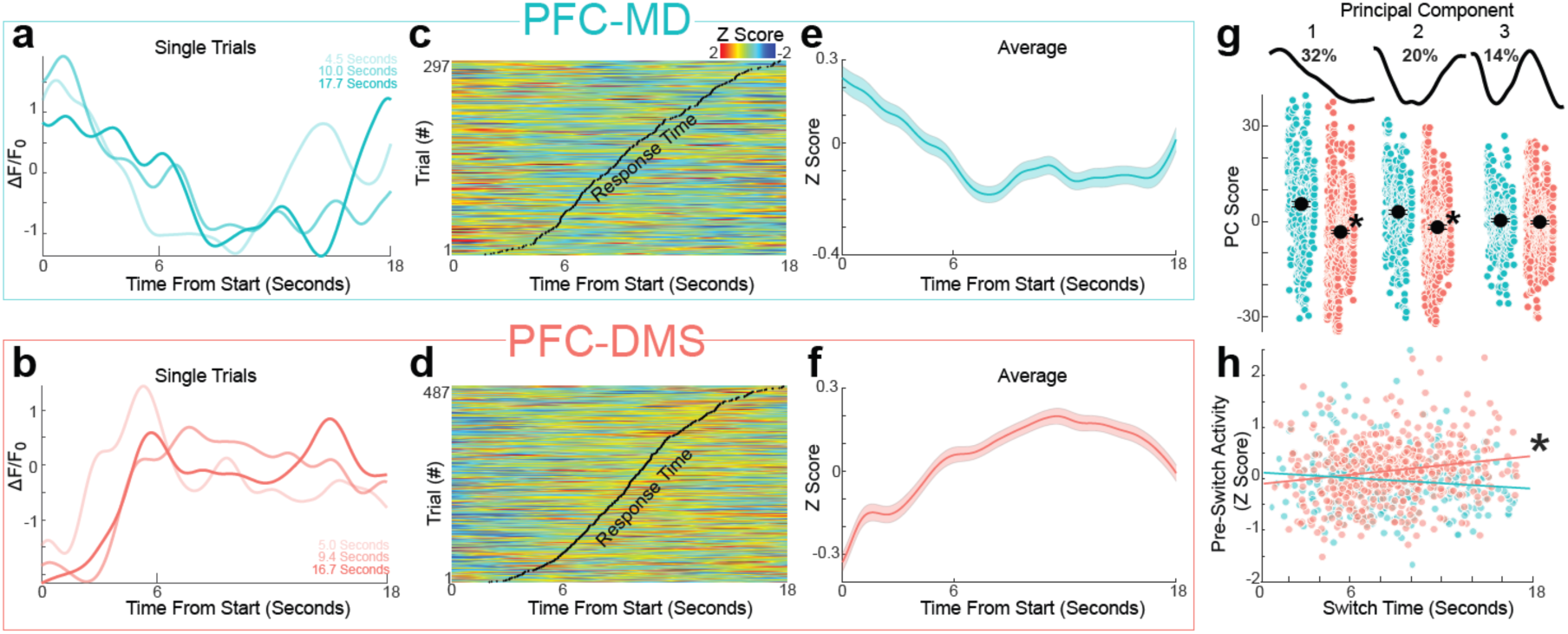
Prefrontal projections have distinct time-related ramping. a) PFC-MD activity (in aqua) and b) PFC-DMS (in rose) from 3 random single trials during interval timing; different shades of aqua and rose represent different response times, shown in the key. c) Z-scored activity from all single trials from all mice during interval timing for PFC-MD projections, which tended to ramp down over the interval. Black dots indicate response times. d) Z-scored activity from all single trials from all mice during interval timing for PFC-DMS projections, which tended to ramp up until the switch. Black dots indicate response times. e) Average PFC-MD activity was markedly distinct from average f) PFC-DMS activity, with PFC-MD ramping down over the interval and PFC-DMS projections ramping up. g) Principal component analysis (PCA) identified principal component 1 (PC1) as time-related ramping from prefrontal projection activity, explaining 32% of variance. PC1 and PC2 were different between PFC-MD (aqua) and PFC-DMS (rose). \**p* < 0.05. h) Pre-switch-response activity in PFC-DMS projections (rose; −2 to −1 seconds prior to the switch) predicted switch response time, whereas pre-switch response activity in PFC-MD projections (aqua) did not. Data from 297 trials from 7 mice for PFC-MD projections, and 487 trials from 9 mice for PFC-DMS projections. \**p* < 0.05.

To quantify differences in PFC-MD and PFC-DMS dynamics during interval timing using data-driven techniques, we turned again to PCA, which we and others have used extensively to capture patterns of neuronal activity^41–44^. We computed PCA from activity on all trials and all mice from both PFC-MD and PFC-DMS projections. Across mice, we found that the most common pattern (PC1) exhibited time-related ramping activity and explained 32% of variance (Fig 3g), which was remarkably convergent with ramping dynamics identified by distinct recording techniques from EEG and in neuronal ensemble recordings (Fig 1f & 1m). PC2 explained 20% of variance, and PC3 explained 14% of variance (Fig 3g).

Remarkably, PC1 was different between PFC-MD and PFC-DMS projections (PC1 scores of activity from all trials in all mice; 5.4 ± 0.9 vs −3.3 ± 0.6; linear mixed-effect model accounting for mouse-specific variance; F_(1,782)_ = 6.1; *p* = 0.01; Fig 3g). PC2 was also distinct between PFC-MD and PFC-DMS projections (3.0 ± 0.7 vs −1.8 ± 0.5; F_(1,782)_ = 4.8, *p* = 0.03), while PC3 was not. These data help delineate the diversity of prefrontal neurons and further demonstrate that PFC-MD and PFC-DMS activity had distinct time-related ramping dynamics.

During interval timing, response time indicates a participant’s internal estimate of time, because the participant is not guided by external cues and is only influenced by temporal control of action^22^. In our mouse-optimized interval timing task, the moment when mice depart the short nosepoke after determining that no reward has been delivered at∼6 seconds reflects the mouse’s internal estimate of elapsed time^27,32^. Although PFC-MD and PFC-DMS activity was not differentially modulated by nosepoke entries (Fig S1), PFC-MD activity ramped down to ∼6 seconds, while PFC-DMS activity appeared to ramp up to the switch response time (Fig 3c–d). To quantify this pattern, we examined pre-switch activity in PFC-DMS projections (average activity from −2 to −1 second prior to the switch, to exclude pre-switch motor activity) and found that pre-switch PFC-DMS activity significantly predicted switch response time (r = 0.17, *p* = 0.0001), whereas pre-switch PFC-MD activity did not (r = −0.11, *p* = 0.06). Critically, pre-switch activity had a distinct relationship with response times for PFC-DMS compared to PFC-MD projections (linear mixed-effect model accounting for mouse-specific variance on response times: pre-switch activity: F_(1,780)_ = 5.3, *p* = 0.02; PFC-DMS vs PFC-MD: F_(1,780)_ = 0.8, *p* = 0.37; interaction: F_(1,780)_ = 9.3, *p* = 0.002). We did not observe other relationships between response times and either starting activity or slope of activity over the interval (Fig S2). These data are particularly impressive given that PFC-DMS fiber photometry integrates bulk activity from populations of neurons expressing DIO-GCaMP6s. Together, these results reveal that while PFC-MD activity ramped down to ∼6 seconds, PFC-DMS activity ramped up until the switch response, reflecting the animals’ internal estimate of time.

### Inactivating PFC-DMS projections increased response times

To examine causal contributions of prefrontal projections, we harnessed circuit-specific optogenetics in which we expressed Archaerhodopsin (Arch) in the PFC and targeted fiber optics to the MD and DMS for terminal inhibition (Fig 4a-c, g). We compared behavior on trials with laser-induced inhibition of prefrontal projections (50% of trials) vs control trials with the laser off (50% of trials). PFC-MD projections and PFC-MD projections were inactivated separately. The experimental cohort consisted of 10 mice: 2 with optical fibers implanted in both the MD and DMS, and 4 with fibers targeting either the MD or the DMS. A separate control group of 4 mice received control EGFP-expressing viruses (lacking opsins) and had fibers implanted in both regions.

**Figure 4:**
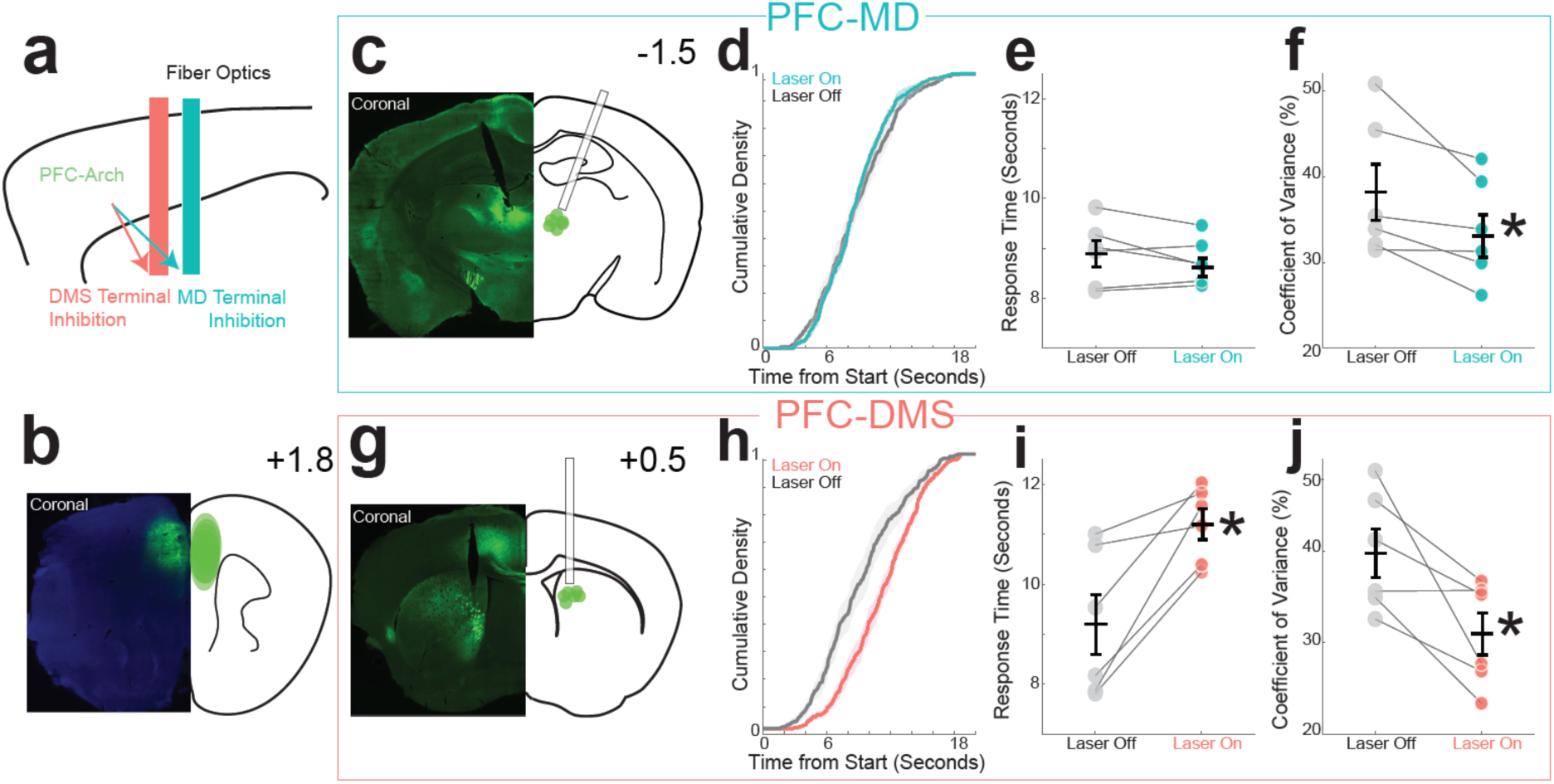
Optogenetic inactivation of prefrontal projections. a) Strategy for optogenetic terminal inhibition of PFC-MD projections (aqua) and PFC-DMS projections (rose) by b) expressing Archaerhodopsin (Arch) into the PFC. Top right number indicates anterior-posterior (AP) coordinate. Arch virus expressions from each mouse shown with a green ellipse. c) We implanted fiber optics bilaterally into the MD. Top right number indicates AP coordinate. Each fiber optic location with Arch virus is shown with a green dot. d) On trials with PFC-MD projections inhibited, we observed little change in the cumulative density of response times. e) There was no change in the average response time, but there was f) a decrease in the response time CV. g) We implanted fiber optics bilaterally into the DMS. Top right number indicates AP coordinate. Each fiber optic location with Arch virus is shown with a green dot. h) On trials with PFC-DMS projections inhibited, we observed marked changes in the cumulative density of response times, with i) an increase in the average response time, and j) a decrease in the response time CV. Data from 10 mice: 2 with fiber optics in both the MD and the DMS, 4 with fiber optics in the MD, and 4 with fiber optics in the DMS. \**p* < 0.05.

We found that inactivating PFC-MD projections did not affect response times (8.9 ± 0.3 seconds vs 8.7 ± 0.02 seconds with PFC-MD inhibited; t_(5)_ = 1.2, *p* = 0.30; Fig 4c–e) but decreased response time CV (38 ± 3% vs 34 ± 2% with PFC-MD inhibited; t_(5)_ = 2.7, *p* = 0.04; Fig 4f). By contrast, inactivating PFC-DMS projections increased response times (9.2 ± 0.6 seconds vs 11.2 ± 0.3 seconds with PFC-DMS inhibition; t_(5)_ = 4.1, *p* = 0.01; Fig 4g–i), providing evidence that PFC-DMS projections contribute to the temporal control of action. Inactivating PFC-DMS projections also decreased response time CV (40 ± 3% vs 31 ± 2% with PFC-DMS inhibition; t_(5)_ = 3.0, *p* = 0.03; Fig 4j). Critically, there were no reliable effects in control mice (Fig S3), and we did not find any evidence that PFC-DMS inactivation affected the number of rewards, total number of nosepokes, or traversal times between the short and long nosepoke ports (Fig S4)^27^. Taken together, these data demonstrate that PFC-MD and PFC-DMS projections played distinct roles in interval timing, with PFC-DMS projections controlling estimates of time.

### Prefrontal projections have distinct transcriptional features

Our work thus far suggests that PFC-MD and PFC-DMS projections have distinct activity patterns and effects on interval timing. To map these differences to transcriptional profiles, we harnessed snRNA-seq using H2B-TRAP mice with retrograde AAV Cre virus injected in either the MD or the DMS to label nuclei in the PFC with mCherry. We physically dissected prefrontal projections by collecting tissue from anterior cingulate, prelimbic, and infralimbic regions of the mouse PFC. Subsequently, we prepared single-nuclei suspensions and performed fluorescence-activated nuclei sorting to collect mCherry-labeled PFC-MD (22 mice) or PFC-DMS (20 mice) projecting nuclei (Fig 5a). After sequencing, we performed read alignment to the mouse genome and applied quality control filters to the data before downstream analysis (Fig S5a).

**Figure 5:**
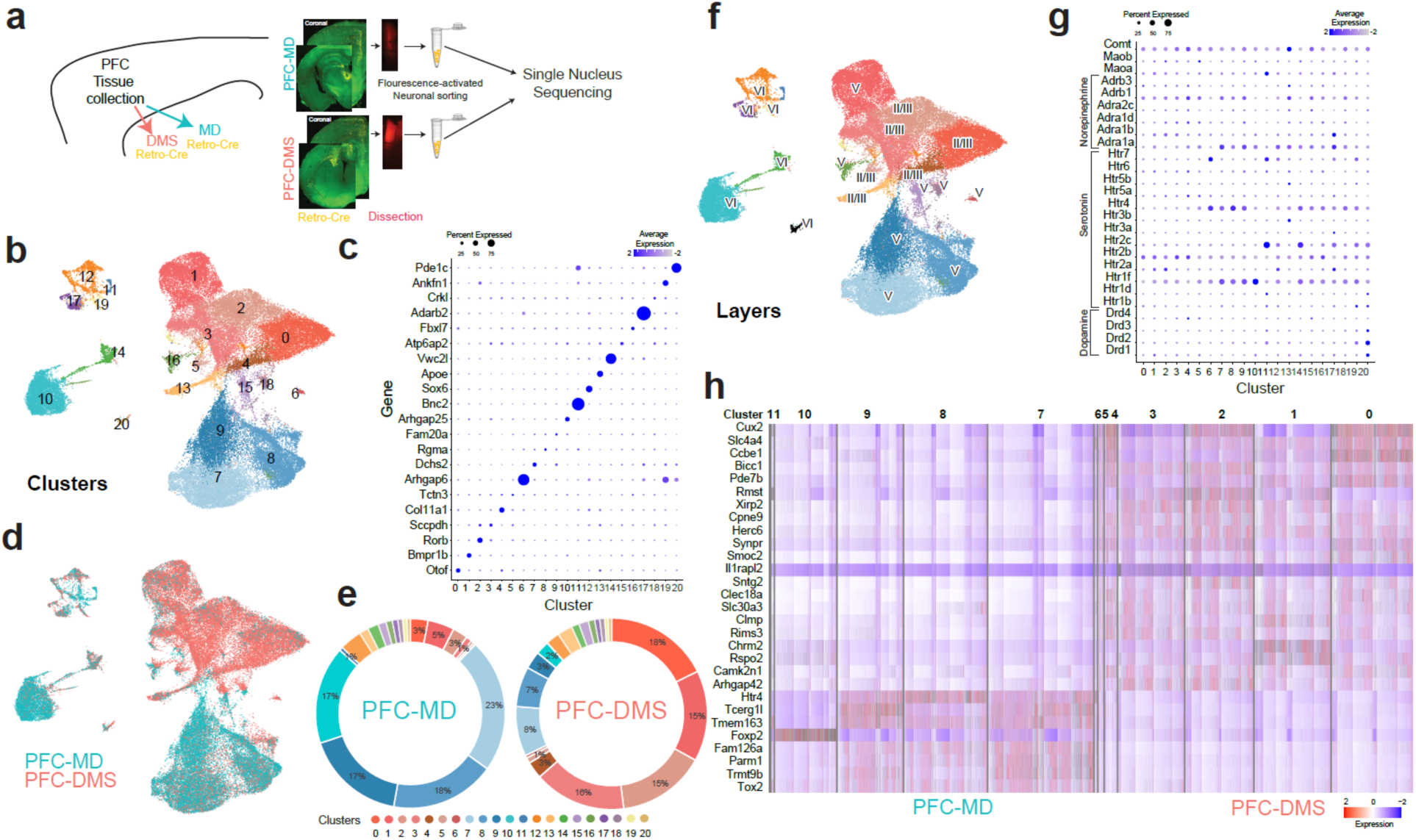
PFC-MD and PFC-DMS have distinct transcriptional profiles. a) Schematic of the experimental design (PFC-MD: 22 mice (12 female); PFC-DMS: 20 mice (11 female). b) Uniform manifold approximation and projection (UMAP) plot shows PFC-MD and PFC-DMS neurons classified into 21 clusters. c) Dotplot of gene markers of each cluster. d) Feature plot showing the distinct expression patterns of PFC-MD (in aqua) and PFC-DMS (in rose). e) The percentage of each cluster in PFC-MD and PFC-DMS projections. See Table 2 for more information. f) Cortical layer assignments for PFC-MD and PFC-DMS clusters. g) Expression of receptors for neuromodulators such as dopamine, serotonin, and norepinephrine, as well as receptors for catechol-o-methyl transferase (Comt) and monoamine oxidase B (MAOb) and A (MAOa). h) Top genes with high and specific expression in PFC-MD enrichment clusters 7–11 and in PFC-DMS clusters 1–6.

We identified 21 transcriptionally distinct prefrontal neuronal clusters from 66,433 neurons, 28,998 of which were collected from PFC-MD and 37,435 that were collected from PFC-DMS projections (Fig 5b–c). As expected for glutamatergic prefrontal projections, each of these 21 neuronal clusters were positive for at least one canonical excitatory marker (i.e., *Slc17a6, Slc17a7*, or *Camk2a*) and lacked strong expression from GABAergic markers such as *Slc32a1* (Fig S5b).

Next, we examined the distinction between the transcriptional profiles of PFC-MD and PFC-DMS projection neurons. We observed enrichment of PFC-MD neurons in clusters 7–11 (defined as the PFC-MD enrichment group; Fig 5d–e) and enrichment of PFC-DMS neurons in clusters 0–6 (defined as the PFC-DMS enrichment group). Neurons in clusters 12–21 were equally distributed between PFC-MD and PFC-DMS neurons. We analyzed the expression pattern of cortical layer-specific marker genes within our dataset. Clusters 0, 2, and 4 clearly showed enrichment of *Cux2* and *Cux1*, which are marker genes for Layer II/III cortical neurons^45,46^. Clusters 6–10 and 20 were enriched for *Bcl11b*, while cluster 14 was enriched for *Etv1*, both of which are enriched in layer V^47–49^. Clusters 10 and 20 were enriched for Layer VI gene markers *Syt 6* (Fig S5d–e)^50,51^. To ensure statistically informed classification, we calculated layer-specific expression scores (as detailed in the Methods) and used these scores to assign each cluster to a cortical layer (Fig 5f, Fig S5c). For PFC-MD enrichment groups, clusters 7–9 were enriched for Layer V, and cluster 10 was enriched for Layer VI; by contrast, for PFC-DMS enrichment groups, clusters 0, 2, 3, 4, and 5 were enriched for Layer II/III, while clusters 1 and 6 were enriched for Layer V. Cluster 11 showed co-expression of *Cux2*, which marks Layer II/III, and *Bcl11b*, which marks Layer V. In combination with light sheet microscopy (Fig 2a), these data provide new evidence that PFC-MD projection neurons are transcriptionally distinct from PFC-DMS projection neurons, with PFC-MD projection neurons localized to deeper layers and PFC-DMS projection neurons more broadly distributed across superficial and deeper layers. Sex differences are described in Fig S6.

### Enriched genes in prefrontal projections

Prefrontal function and interval timing can be powerfully affected by neuromodulators such as dopamine, serotonin, and norepinephrine^52,53^. We detected the expression of several gene markers of common neuromodulator receptors in all 21 clusters (Fig 5g). Noradrenergic receptor expression was low (<25%) but was stronger in PFC-DMS enrichment groups compared to those of PFC-MD (t = 3.17, *p* = 0.0016). Serotonergic receptors were more expressed in PFC-MD enrichment groups (t = 21.95, *p* < 2.2^e-16^), with *Htr1f* highly enriched in clusters 7–10, *Htr4* highly enriched in clusters 6–9, and *Htr2c* highly enriched in cluster 11 (>30%). Dopaminergic receptors had low expression in both projections (<1%).

We also analyzed genes enriched in the PFC-MD and PFC-DMS clusters and identified highly specific genes based on the criteria described in the Methods section (Figure 5h). *Cux2* was a notable gene in the PFC-DMS clusters; it plays a critical role in cognitive functions and is associated with cognitive disorders^54,55^. It is also a marker for upper cortical layers and supports our previous observation that PFC-DMS neurons are enriched in upper cortical layers. Another gene that was enriched in the PFC-DMS clusters was *Camk2n1*, which regulates calcium signaling^56^. Within the PFC-MD clusters, two genes that were enriched included *Htr4*, which marks serotonin receptors, and *Foxp2*, which has been associated broadly with language and cognitive function^57,58^. Together, these findings suggest that a set of functionally relevant genes underlies the distinctions between PFC-MD and PFC-DMS pathways.

## Discussion

We harnessed circuit-specific techniques to elucidate the diversity of prefrontal activity during temporal control of action. We provide novel evidence that 1) PFC-MD and PFC-DMS activity have distinct patterns of time-dependent ramping, with PFC-DMS activity predicting temporal control of action; 2) inactivating PFC-DMS, but not PFC-MD, influenced internal estimates of time; and 3) PFC-MD and PFC-DMS projections had distinct transcriptional profiles, with PFC-DMS enriched for the Layer II/III marker *Cux2* and the calcium regulator *Camk2n1*, and PFC-MD enriched for the serotonin receptor *Htr4* and the transcription factor *Foxp2*. Taken together, these data provide new insight into the diversity of prefrontal function and will advance new biomarkers and therapeutic interventions targeting prefrontal circuits.

Prefrontal networks have a wide range of inputs and outputs^7,8^; thus, nearly every behavioral paradigm produces a diversity of prefrontal activity^59^. Three past studies have shown that prefrontal projections to the striatum are distinct from projections to the thalamus. Otis et al. (2017) investigated projections to the ventral striatum and paraventricular thalamus and found opposing patterns of activity around reward-predictive cues^10^. De Kloet (2021) studied prefrontal projections to the mediodorsal thalamus and the dorsomedial striatum during a simple reaction-time task and found bidirectional patterns of PFC-MD and PFC-DMS activity^11^. Wilhem et al. (2023) found that PFC-DMS projections encode and control working memory maintenance activity^12^. Each of these paradigms involves temporal control of action. In the current study we isolated temporal control of action while recording from, manipulating, and transcriptomically characterizing PFC-MD and PFC-DMS projections.

Over any temporal interval, PFC neurons exhibit both increasing and decreasing time-dependent ramping activity across tasks and species^17,18,42,43^. In prior studies^10,12^, PFC-MD and PFC-DMS projections demonstrate functionally distinct patterns of activity. An advance of this study is that we identified that these opposing patterns are characterized by time-dependent ramping activity, a well-described temporal code^23,60^. Ramping activity is computationally consistent with drift-diffusion models that integrate the accumulation of temporal evidence^25,26^. While prior studies have consistently shown that prefrontal networks robustly encode time and exhibit time-dependent ramping activity^17,18,24,30,42–44^, our study breaks new ground by revealing that PFC-DMS activity ramps up to response times, suggesting that PFC-DMS activity represents animals’ internal estimates of time. This idea is supported by our data that disrupting PFC-DMS projections prolongs response time, underscoring the essential role of PFC-DMS projections in temporal control of action. These data, combined with our previous findings that striatal ramping requires prefrontal function and that stimulating PFC-DMS projections can rescue timing deficits caused by prefrontal inactivation^17,18,61^, strongly suggest that temporal signals in the striatum originate from the PFC.

We also found that PFC-MD projections ramped down over the timed interval and were prominently modulated by reward. Inactivating PFC-MD projections did not affect response time, suggesting a difference between ramping activity in PFC-MD vs. PFC-DMS projections. Indeed, PFC-MD projections ramped down throughout the interval before increasing at reward, while PFC-DMS neurons ramped up to response time. These data indicate that the direction of cortical ramping conveys distinct computational information.

Fascinatingly, inactivating both PFC-MD and PFC-DMS decreased timing variability. In prior studies, MD inactivation increased both response time and variability^18,62^. Furthermore, the MD integrates feedforward basal ganglia inputs downstream from PFC-DMS projections. It may be that PFC-MD projections regulate timing precision, but PFC-DMS projections regulate timing accuracy, and effects on timing precision are secondary. Future studies that record from mediodorsal thalamus with and without prefrontal inactivation may clarify how thalamocortical networks contribute to timing variability and why PFC projection inactivation decreases timing variability.

We also extend past work^10–12^ by linking projection-specific prefrontal activity to distinct molecular motifs. Several previous studies have investigated the molecular profiles of prefrontal projections, and some of these studies revealed that projections to the striatum and the mediodorsal thalamus are enriched in superficial vs deep cortical layers^8,63^. Single-cell transcriptomic approaches have provided new information about prefrontal cell types and have contributed to understanding the molecular architecture of the PFC^64–68^. Our study adds new information about PFC-MD and PFC-DMS projections and the functional relevance of prefrontal transcriptional profiles. For instance, neither PFC-MD nor PFC-DMS projections strongly expressed dopamine or norepinephrine receptors, but PFC-MD projections strongly expressed *5HT4* serotonin receptors, which might be helpful for therapies for human diseases that disrupt PFC^69^. Moreover, laminar markers such as *Cux2* exhibited distinct expression patterns in PFC-DMS projections compared to PFC-MD projections^45^. Additionally, *Foxp2*, which is highly expressed in PFC-MD projections, plays a fascinating role in language, regulates dopamine receptors, and is specifically attenuated in prefrontal regions of patients with Parkinson’s disease^57,58,70^. Despite the challenges of linking PFC activity to transcriptional profiles, our data offer insight into the potential role of distinct gene expression patterns associated with the temporal control of action.

Our approach has several limitations. First, our fiber photometry techniques record aggregate activity of prefrontal projections. Consequently, further dissection of PFC based on activity patterns may be insightful. Second, it is difficult to delineate the contribution of PFC-MD and PFC-DMS projections to prefrontal neuronal ensemble averages or to cortical event-related potentials (ERPs); future studies might combine prefrontal neurophysiology with circuit-specific techniques. Third, in this study we did not stimulate PFC-MD and PFC-DMS projections, although our prior work found no effect of PFC-DMS stimulation except when prefrontal circuits were dysfunctional^61^. Lastly, we did not link specific transcriptional profiles with neuronal activity. Future work will combine manipulation of specific genes with optogenetics and neuronal recordings to determine precisely what information prefrontal projections contribute to downstream brain areas during timing.

In summary, our work shows that PFC-MD and PFC-DMS projections have distinct patterns of ramping activity. Inactivating PFC-MD and PFC-DMS projections affected timing variability, while only inactivating PFC-DMS projections affected response time. Finally, transcriptional profiling revealed that PFC-MD and PFC-DMS exhibited distinct patterns of gene expression. Taken together, our data on prefrontal projections help us understand the diversity of PFC activity during cognitive functions.

## Methods and Materials

### Human participants

A total of 40 healthy participants under 50 years old were recruited to perform an interval timing task. Of these participants, 32 were female (median age: 29.1 years; interquartile range (IQR): 16.2 years), and 12 were male (median age: 27.4 years; IQR: 15.5 years). All participants were recruited under University of Iowa Institutional Review Board #201707828. Behavioral data from some of the same participants have been reported previously^31^.

### Human interval timing

We adapted an interval timing switch task that has previously been used in rodents and humans^32,36,37^. This task required participants to estimate a short time interval (2 seconds) before the end of a longer time interval (3 seconds) and indicate their time estimate by switching from the left-arrow key to the right-arrow key (Fig 1a). The task was performed on a Dell XPS workstation with a 19-inch monitor. Task-specific audio was played through Dell Rev A01 speakers positioned on either side of the monitor, and responses were made with the left and right index fingers on a standard QWERTY USB-keyboard. There was an equal number of two types of trials: short interval trials and long interval trials; participants were not told the interval length at the start of each trial. Participants were trained using a three-phase approach. Phase 1) Participants observed short and long intervals by watching a box appear on the screen for either a short or long interval of time; the displayed box was accompanied by the word “short” or “long,” respectively. Phase 2) Participants observed 10 trials during which the box appeared for either a short or long interval and verbally stated whether the interval was short or long. Participants received immediate verbal feedback from the experimenter; they were required to achieve 80% accuracy to pass this stage. Phase 3) Participants completed a single 3-minute training block, during which visual cues on the screen indicated accuracy, but no points were earned. Testing consisted of four 4-minute blocks of trials, with ∼60 trials per block.

All interval timing trials started with a white fixation cross displayed on a blank gray screen; the appearance of the white fixation cross cued participants to press and hold the left arrow key, which caused an empty box outlined in green to appear on the left side of the screen. In short interval trials, the outlined box switched to a solid green box at 2 seconds, and the participant received 1 point as reinforcement. If the participant responded by switching keys before 2 seconds had elapsed, they did not earn a point. In long interval trials, the outlined box switched to a solid green box at 3 seconds. Participants were asked to respond after 2 seconds but before 3 seconds had elapsed by switching from pressing and holding the left-arrow key to pressing and holding the right-arrow key. This response caused the box outlined in green to switch from the left side of the screen to the right side; then at 3 seconds, the outlined box on the right side switched to a solid green box. If the participant correctly responded before 3 seconds, the participant received 1 point. If the participant failed to switch keys before 3 seconds, they did not earn a point. The decision to switch is guided explicitly by an internal estimate of time. Only the data from trials in which a “switch” occurred were analyzed.

### Human EEG

During EEG recordings, each participant sat in a quiet room in front of a computer monitor to participate in behavioral assays. All task stimuli were presented using the Psychtoolbox-3 functions^31,71^ in MATLAB. EEG recordings were performed according to methods described in detail previously^72,73^. Briefly, we used a 64-channel EEG actiCAP (Brain Products GmbH, Gilching, Germany) with a 0.1-Hz high-pass filter and a sampling frequency of 500 Hz. We used electrode Pz as the online reference and electrode site Fpz for the ground. EEG activity was referenced according to the procedures described in Singh et al. 2023^72^. An additional channel was recorded at the mid-inion region (Iz), and we removed unreliable FT9, FT10, TP9, and TP10 channels, resulting in 59 channels for pre- and post-processing. Data were epoched around the cues from −1 to 2.5 seconds peri-cue. Bad channels and bad epochs were identified using a combination of the Fully Automated Statistical Thresholding for EEG artifact Rejection (FASTER) algorithm that rejects artifacts with greater than ± 3 z-scores on key metrics and the pop_rejchan function from EEGLAB Version 14. Subsequently, bad channels were interpolated, with the exception of the midfrontal “Cz” channel. Eye blink contaminants were removed following independent component analysis (ICA). ERPs were low-pass filtered at 20 Hz for analyses. PCA was performed across participants and electrodes.

### Rodents

A total of 92 mice were used across all mice experiments (overview in Table 1). For electrophysiology recording, circuit mapping study, and fiber photometry and optogenetics experiments, 50 wild-type C57BL/6J mice were obtained from The Jackson Laboratory (Jax #000664; Bar Harbor, ME). For snRNA-seq experiments, 42 transgenic H2B-TRAP (±) mice were generated by crossing H2B-TRAP mice (Dr. Jon Resch’s laboratory at the University of Iowa, Jax #029789)^74^ with C57BL/6J mice from The Jackson Laboratory. All experimental mice were 3–4 months of age at the time of surgery and were group housed under a 12-hour light-dark cycle in a temperature- and humidity-controlled room with *ad libitum* access to standard laboratory rodent chow and water. All behavioral mice were singly housed following surgery with *ad libitum* access to food and water, until starting food restriction at ∼85% of free-fed weight 1 week after surgery. All experiments were performed during the animals’ light period. All procedures were approved by the Institutional Animal Care and Use Committee (IACUC) at the University of Iowa and performed in accordance with applicable guidelines and regulations, Protocol #0062039.

**Table 1:**
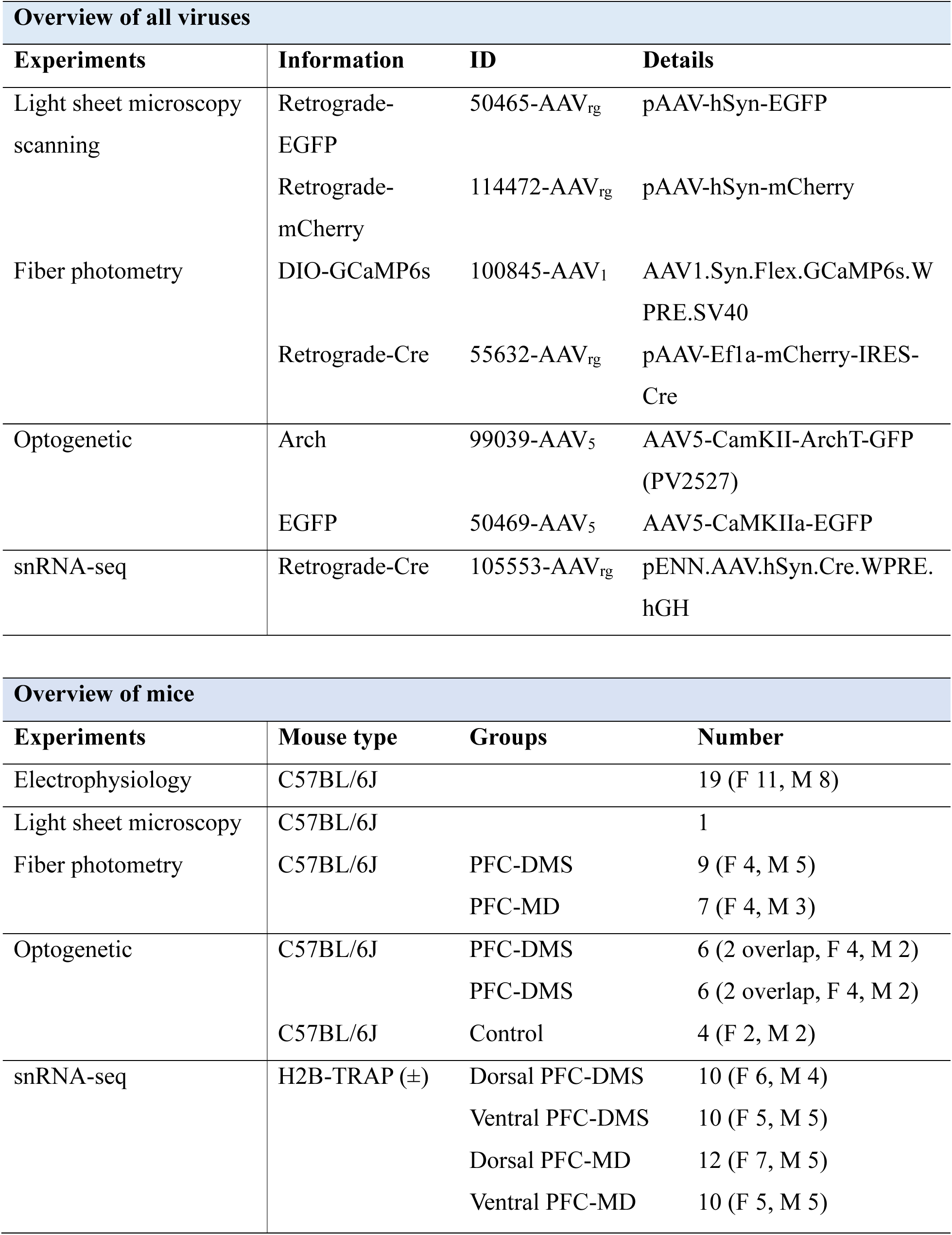
Key resource table.

**Table 2:** Cluster expression profiles for each prefrontal projection.

| **Cluster** | **PFC-MD** | **PFC-DMS** |
| --- | --- | --- |
| **0** | 17.70% | 3.02% |
| **1** | 15.36% | 4.54% |
| **2** | 14.91% | 2.58% |
| **3** | 15.59% | 1.03% |
| **4** | 2.86% | 0.41% |
| **5** | 0.94% | 0.37% |
| **6** | 0.46% | 0.08% |
| **7** | 8.40% | 22.93% |
| **8** | 6.77% | 17.88% |
| **9** | 2.98% | 17.10% |
| **10** | 2.29% | 16.71% |
| **11** | 0.05% | 0.63% |
| **12** | 2.29% | 3.78% |
| **13** | 2.54% | 1.57% |
| **14** | 1.31% | 1.86% |
| **15** | 1.62% | 1.38% |
| **16** | 1.08% | 1.08% |
| **17** | 0.83% | 0.93% |
| **18** | 0.85% | 0.81% |
| **19** | 0.59% | 0.76% |
| **20** | 0.57% | 0.56% |

### Viral vectors

All viruses used in this study are shown in Table 1.

### Surgical procedures

Mice were anesthetized with 1.5%–4.0% isoflurane gas delivered at 120 mL/min (SomnoSuite, Kent Scientific, Torrington, CT). Mice were placed on a heating pad in a stereotaxic frame, and the scalp was retracted to expose the skull. Depending on the experiment, craniotomies were drilled above the PFC and the mediodorsal thalamus (MD) or the dorsomedial striatum (DMS), according to the coordinates stated below. Viruses (described in the key source table, Table 1) were infused over 10 minutes (0.05 µl/min; Legato 130 Syringe Pump, kd Scientific, Holliston, MA), with a 10-minute wait period before removing the needle. During electrophysiology, fiber photometry, and optogenetics experiments, we attached at least three stainless steel screws to the skull to stabilize head cap assemblies, which were sealed with cyanoacrylate (“SloZap,” Pacer Technologies, Rancho Cucamonga, CA) and accelerated with “ZipKicker” (Pacer Technologies) and methyl methacrylate (AM Systems, Port Angeles, WA). Following all surgical procedures, mice were allowed at least 1 week to recover before starting food restriction and interval timing training.

For mouse electrophysiology, a craniotomy was drilled above the PFC (AP +1.8, ML ±0.3), and a 4×4 (1 mm^2^) multielectrode recording array (MicroProbes, Gaithersburg, MD) was implanted (DV −1.8). Electrode arrays were counterbalanced left vs right hemisphere across mice. A stainless-steel wire was secured to at least one skull screw to ground the electrode array. See prior work for electrophysiological details^27,75^.

For circuit mapping, two fluorescents retrograde AAV tracers were injected in both the MD (0.5 µl: AP −1.5, ML −1.63, DV −3.8 at 20° lateral angle) and the DMS (0.5 µl: AP +0.5, ML-1.4, DV −2.7) in the same mouse and the same hemisphere. The mouse was allowed to recover for 3 weeks to ensure robust fluorescent expression before tissue collection, described below.

For fiber photometry experiments, the retrograde AAV Cre virus was unilaterally injected in either the MD (0.5 µl: AP −1.5, ML +1.63 or −1.63, DV −3.8 at 20° lateral angle) or the DMS (0.5 µl: AP +0.5, ML+1.4 or −1.4, DV −2.7) in different cohorts of mice. In all mice, a Cre-dependent GCaMP6s virus was injected into the PFC (PFC-MD: AP +1.8, ML +0.45 or −0.45, DV −1.7/-1.8 at 0.5 µl/DV; PFC-DMS: AP +1.8, ML +0.35 or −0.35, DV −1.7/-1.8 at 0.5 µl/DV). A fiber optic cannula (Doric Lenses Inc, Quebec, Canada) was then implanted in the PFC (PFC-DMS: AP +1.8, ML 0.35 or −0.35, DV −1.8; PFC-MD: AP +1.8, ML 0.45 or −0.45, DV −1.8).

For optogenetics experiments, Archaerhodopsin (Arch) virus was bilaterally injected in the PFC (AP +1.8, ML±0.4, DV −1.8, 0.5 µl/DV) followed by bilateral fiber optic cannulas (Doric Lenses Inc, Quebec, Canada) implanted in the MD or the DMS, using the same coordinates as the fiber photometry experiments. Each group had 6 mice; 2 mice had fibers in both the MD and the DMS, while 4 mice had fibers in only the MD or the DMS. A separate group of 4 mice were injected with EGFP virus instead of the Arch virus, as controls. In all 4 control mice, fiber optic cannulas were implanted bilaterally in both the MD and the DMS using the same coordinates as above.

In the snRNA-seq experiments, we assessed the transcriptional profiles of both dorsal and ventral PFC populations that project to either the MD or the DMS. To access PFC populations projecting to either the MD or the DMS, cohorts of both male and female H2B-TRAP mice received stereotaxic bilateral injections of retrograde AAV Cre into either the DMS (AP +0.5, ML ±1.4, DV −2.7, 0.5 µl/DV) or the MD (AP −1.5, ML ±1.63, DV −3.8 at 20° lateral angle, 0.5 µl/DV). The details of each group are shown in the key source table (Table 1).

### Mouse interval timing

We used a mouse-optimized operant interval timing task described in detail previously^27,32,36,37^. Briefly, mice were trained in sound-attenuating operant chambers with two front nosepoke ports (“short” and “long”) flanking either side of a food hopper on the front wall and a third nosepoke port located at the center of the back wall. The chamber was positioned below an 8-kHz, 72-dB speaker (Fig 1h; MedAssociates, St. Albans, VT). Mice were 85% food restricted and motivated with 20-mg sucrose pellets (Bio-Serv, Flemington, NJ). Mice were initially trained to receive rewards during fixed-ratio nosepoke response trials. Nosepoke entry and exit were captured by infrared beams. After shaping, mice were trained in the “switch” interval timing task. Mice self-initiated trials at the back nosepoke, after which a tone and nosepoke lights were illuminated simultaneously. Cues were identical on all trial types and lasted the entire trial duration (6 seconds: short trials or 18 seconds: long trials). On 50% of trials, mice were rewarded for a nosepoke after 6 seconds at the designated short nosepoke port; these trials were not analyzed. On the remaining 50% of trials, mice were rewarded for responding first at the short nosepoke port and then switching their response to the long nosepoke port; a reward was delivered for an initial response at the long nosepoke after 18 seconds. Early responses at the short or long nosepoke port were not reinforced. Error trials were rare and included trials where animals responded only at the short or long nosepoke port, which were also not reinforced.

The two primary measures of interval timing performance were: 1) timing accuracy, quantified by the average switch response time of all switch trials performed during an interval timing session; and 2) timing precision, quantified by the switch response time coefficient of variance (CV) of all switch trials performed during an interval timing session, calculated as the standard deviation divided by the average and expressed as a percentage. We also analyzed the number of rewards, the number of responses, and the traversal time from the final short nosepoke to the first long nosepoke during a switch response.

### Light sheet microscopy

Mouse brain was collected and fixed in 4% paraformaldehyde (PFA) at 4 °C overnight. After fixation, the samples were washed and stored in phosphate-buffered saline (PBS; pH 7.4) before undergoing optical clearing using the uDISCO method^76^. In brief, the whole brain was sequentially incubated in tert-butanol solutions of increasing concentrations (30%, 50%, 70%, 80%, 90%, 96%, and 100% v/v) each for 24 hours. The brain was then incubated in dichloromethane (DCM) at room temperature for 45–60 minutes, followed by immersion in BABB-D10 for 2–6 hours, until the sample became optically transparent. Light sheet microscopy was performed using a 3i AxL Cleared Tissue Light Sheet microscope (Intelligent Imaging Innovations, Denver, CO) running Slidebook software. Brains were imaged with a 1x/0.25NA objective at 1 micron voxel size (high axially scanned light-sheet microscopy mode) using 488-nm (GFP) and 561-nm (mCherry) laser light with corresponding 525/50-nm and 605/64-nm emission filters. 3D image tiles were aligned using the built-in “zero shifts” algorithm in Slidebook.

### Electrophysiology

We recorded single prefrontal neuron activity using a multielectrode recording system (Open Ephys, Atlanta, GA) during interval timing. Offline Sorter™ (Plexon, Dallas, TX) was used to remove artifacts and classify single units using PCA and waveform shape: single units were defined as those with a consistent waveform shape, a separable cluster in PC space, and a 2-ms or longer refractory period in the inter-spike interval histogram. Spike activity was calculated using kernel density estimates of firing rates across the interval, binned at 0.25 seconds with a bandwidth of 1. We focused our analysis on interval timing switch trials only. We constructed peri-event time histograms (PETHs) using kernel density estimates and z-score normalization of firing rates between −4 seconds and 22 seconds from trial start. We then used PCA to quantify patterns of activity across trials and mice. To measure time-related ramping over the 18-second interval, we used trial-by-trial GLMs at the individual neuron level in which the response variable was firing rate and the predictor variable was time in the interval, as in our past work^17,27,44^. For each neuron, its time-related slope was derived from the GLM fit of firing rate vs time in the interval, for all trials per neuron. All GLMs were run at a trial-by-trial level to avoid the effects of trial averaging, and p-values were corrected via false-discovery rate ^77^.

### Fiber photometry

Mice were extensively trained in the interval timing switch task described above before acclimation to fiber photometry procedures. Mice were tethered to a unilateral optical patch cord connected to a Fiber Photometry System from Doric Lenses Inc (Québec, Canada) with two excitation wavelengths (405-nm LED isosbestic signal modulation at 208.616 Hz and 470-nm LED Ca^2+^-dependent GCaMP6s signal modulated at 572.205 Hz). Light was emitted, and isosbestic and GCaMP6s fluorescence wavelengths were measured via integrated LED mini cubes and photo detector (iLFMC4, Doric Lenses Inc). Recording sessions lasted 60 minutes to avoid photobleaching artifacts. Changes in GCaMP6s fluorescence was aligned to task events, downsampled to 100 Hz, low-pass filtered at 5 Hz, detrended to compensate for exponential decay and photobleaching, motion corrected via the isosbestic signal, normalized to obtain ΔF/F_0_, and then z-scored for further analysis^78^.

### Optogenetics

Mice were extensively trained in the interval timing switch task described above before acclimation to optogenetic procedures, described previously^20,61,79^. Briefly, mice were tethered to a 520-nm laser diode (Doric Lenses Inc) through a bilateral optical rotary joint and optical patch cords (Doric Lenses Inc). Pathway-specific optogenetic inhibition was randomly delivered during the entire trial (6 or 18 seconds) on 50% of trials. Laser power output was adjusted in separate behavioral sessions to 20 mW at the tip of the optical patch cord. Each experimental session lasted 90 minutes. We focused on interval timing switch trials and compared average switch response time and switch response time CV when no laser light was delivered (Laser Off) vs when laser light was delivered (Laser On) during the entire trial.

### Histology

Mice were sacrificed following the completion of electrophysiology, fiber photometry, and optogenetics experiments. Mice were deeply anesthetized using ketamine (100 mg/kg IP) and xylazine (10 mg/kg IP) and transcardially perfused with 1x PBS followed by 4% PFA. Brains were removed and fixed in 4% PFA overnight and then transferred to a 30% sucrose solution for approximately 48 hours. Fixed brains were sliced into 40-μm coronal sections throughout the PFC, DMS, and MD using a sliding microtome (HM 430, Epredia, Kalamazoo, MI). Sections were collected and mounted directly on positively-charged slides with ProLong Diamond Antifade Mountant with DAPI (P36962, Invitrogen, Waltham, MA). Brain sections were imaged using an Olympus VS120 slide scanning microscope (Olympus, Center Valley, PA).

### Single-nucleus RNA sequencing

After 2 weeks of recovery from viral injection surgeries, mice were sacrificed by rapid decapitation, after removal from the home cage, to limit transcriptional changes due to stress and anesthesia. Each brain was then rapidly extracted and immediately placed in slushed DMEM/F12 (1:1) solution (Thermo Fisher Scientific, Waltham, MA) for approximately 1 minute. Coronal sections containing the PFC were then sliced using a clean razor blade and a 1-mm chilled stainless steel brain matrix (Stoelting, Wood Dale, IL) and immediately placed in chilled RNAprotect solution (QIAGEN, Aarhus, Denmark). Next, the labeled dorsal or ventral region of the PFC region was microdissected under a fluorescent stereoscope (Nikon SMZ800N, Tokyo, Japan) using a 1-mm Heath Curette (Roboz Surgical Instrument Co, Gaithersburg, MD). Dissected tissues were incubated overnight at 4° C in RNAprotect® before being removed and stored dry at −80 °C.

Tissue was then homogenized into a single-nuclei suspension, and nuclei that were positive for mCherry fluorescent labeling were collected using fluorescence-activated nuclei sorting (FANS). Nuclei isolation, FANS, and Chromium Next GEM sequencing library prep protocols were completed, as described in Schwalbe et al. 2024with minor modifications^80^. FANS sorting was completed on a CYTEK Aurora™ cell sorter (CYTEK, Fremont, CA) with a 70-µm nozzle, with assistance from experts in the University of Iowa Flow Cytometry Facility. snRNA-seq libraries were sequenced, targeting 20,000 reads per nucleus, on an Illumina NovaSeq6000 Sequencer (San Diego, CA) operated by specialists at the University of Iowa Institute of Human Genetics Genomics Division. After sequencing, demultiplexed sequencing reads were mapped to the mm10-2020-A mouse genome using the Cell Ranger (version 7.2.0) pipeline. Subsequently, all sequencing runs were processed through the CellBender (version 0.2.0) *remove-background* function to filter out reads from ambient RNA and random barcode swapping^81^. In total, two sequencing runs with four samples each were performed, to collect data from both dorsal and ventral PFC populations projecting to either the DMS or the MD from both male and female mice.

For snRNA-seq analyses, we loaded the count matrixes from each run into SeuratObject (version 5.0.2) in R (version 4.4.1) and performed quality control analysis for all sequencing runs. Any nuclei that failed one or more of the following quality control filters were removed from the dataset: >250 unique genes (nFeature); >500 unique molecules (nCount); <10% mitochondrial gene expression; and log10 (nFeature/nCount) > 0.8. Additionally, any genes with 0 reads across all nuclei were removed, and any genes expressed in <10 nuclei were removed to improve analysis processing speeds. Following quality control, all sequencing runs were merged into a single Seurat object and analyzed through the standard Seurat analysis pipeline utilizing the *SCTransform* function (vst.flavor set to “v2”) with mitochondrial gene expression regression. This command normalized our raw counts and identified and scaled the top 3000 variable features. We utilized the Seurat *IntegrateLayers()* function to complete an reciprocal principal component analysis (RPCA)-based batch correction to integrate our 8 samples^82^. After integration, the Seurat *FindNeighbors()*, *FindClusters()*, and *RunUMAP()* functions were used with default settings to cluster and plot our data. To identify differentially-expressed genes, we performed non-parametric Wilcoxon rank sum tests with Bonferroni correction for multiple comparisons. Only genes found to have a corrected p-value of <0.01 and a log-fold change in expression >0.2 were considered to be differentially expressed. Clusters were analyzed for the expression of known cell-type marker genes. Non-neuronal populations, such as astrocytes and oligodendrocytes, were excluded based on marker expression. Clusters that were positive for multiple cell-type markers were labeled as “multiplet” clusters and removed from the dataset. If any “multiplet” clusters were removed, the post-merging analysis above was repeated until no additional “multiplet” clusters were detected.

To determine if a cluster was enriched with prefrontal neurons that project to either the DMS or the MD, we calculated the proportion of total cells from each projection in each cluster. For each cluster, we compared the proportion of total PFC-DMS and PFC-MD neurons. If the proportion of PFC-DMS neurons was more than double that of PFC-MD neurons, we defined that cluster as PFC-DMS-enriched, and vice versa to determine the proportion of PFC-MD neurons. Clusters 0–6 were defined as PFC-DMS-enriched clusters, and clusters 7–11 were defined as PFC-MD-enriched clusters. This definition was applied to all analyses comparing PFC-DMS and PFC-MD neurons.

To classify each cluster into different layers, we applied a set of layer-specific markers based on prior work. The selected markers included: *Cux1* and *Cux2* for Layer II/III; *Bcl11b*, *Fezf2*, *Pou3f1*, and *Etv1* for LayerV^83^; and *Foxp2*, *Sox6*, *Syt6*, and *Tle4* for layerVI^51,64,65,70^. Layer scores were calculated using the *AddModuleScore()* function in Seurat, which allowed us to assign a likely cortical layer identity to each cluster.

To assess neurotransmitter receptor expression across enrichment clusters, we utilized *Drd1*, *Drd2*, *Drd3*, and *Drd4* to identify dopaminergic receptors; *Adra1a*, *Adra1b*, *Adra1d*, *Adra2c*, *Adrb1*, and *Adrb3* for noradrenergic receptors; and *Htr1b*, *Htr1d*, *Htr1f*, *Htr2a*, *Htr2b*, *Htr2c*, *Htr3a*, *Htr3b*, *Htr4*, *Htr5a*, *Htr5b*, *Htr6*, and *Htr7* for serotonergic receptors^84^. We then quantified the expression levels of each receptor in the PFC-DMS and PFC-MD enrichment clusters and compared them using the t.test() function in R.

Finally, we performed differential gene expression analysis comparing PFC-DMS- and PFC-MD-enriched clusters using the *Findmarkers()* in Seurat. Genes were considered differentially expressed if they had a Bonferroni-corrected p-value of <0.01 and a log-fold change in expression >0.2. From these, we identified highly specific marker genes for each group by selecting genes expressed by ≥15% in the enriched group and ≤5% in the opposing group.

### Data and statistics

All data were analyzed with custom code written in MATLAB (R2023a) and R, available at http://narayanan.lab.uiowa.edu/article/datasets. Differences between fiber photometry signals were analyzed at the mouse level using t-tests. Trial-by-trial differences across animals were analyzed using linear mixed-effects models (*fitglme* in MATLAB) including mouse-specific variance. Differences in behavior between optogenetic conditions were analyzed using a paired t-test. Expression levels of each gene between PFC-DMS and PFC-MD enrichment clusters were compared using a t-test. All data are available at http://narayanan.lab.uiowa.edu/article/datasets.

## Supplementary Figures

**Figure S1:**
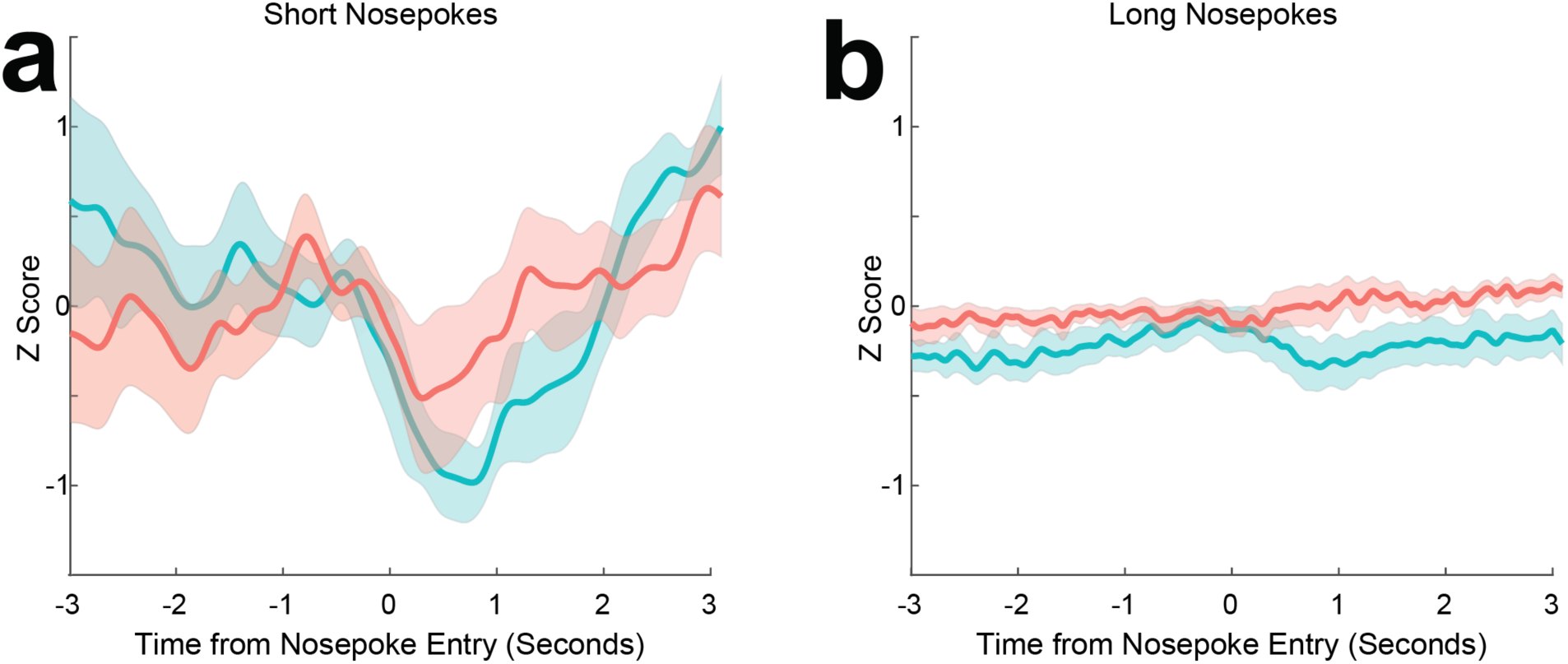
Nosepoke-related activity in prefrontal projections. Calcium activity in PFC-MD (aqua) and PFC-DMS (rose) projections around a) short and b) long nosepoke port entry.

**Figure S2:**
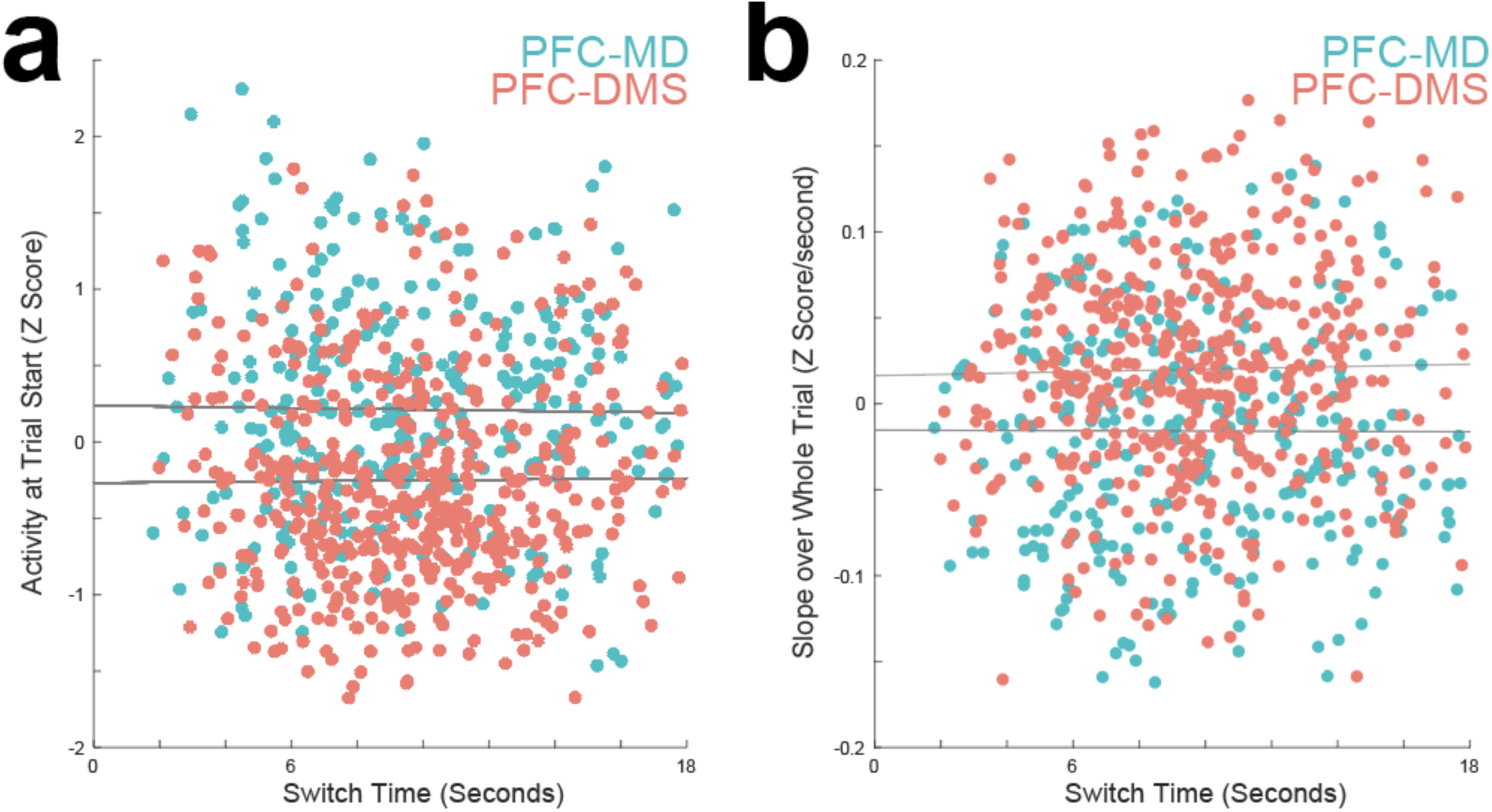
Correlations with switch time. a) Z-scored activity in PFC-MD (aqua) and PFC-DMS (rose) projections for 0–1 second after trial start, plotted as a function of switch response time. b) Linear slope of activity over the 18 second interval for PFC-MD and PFC-DMS as a function of switch response time. Same mice as in Figure 3.

**Figure S3:**
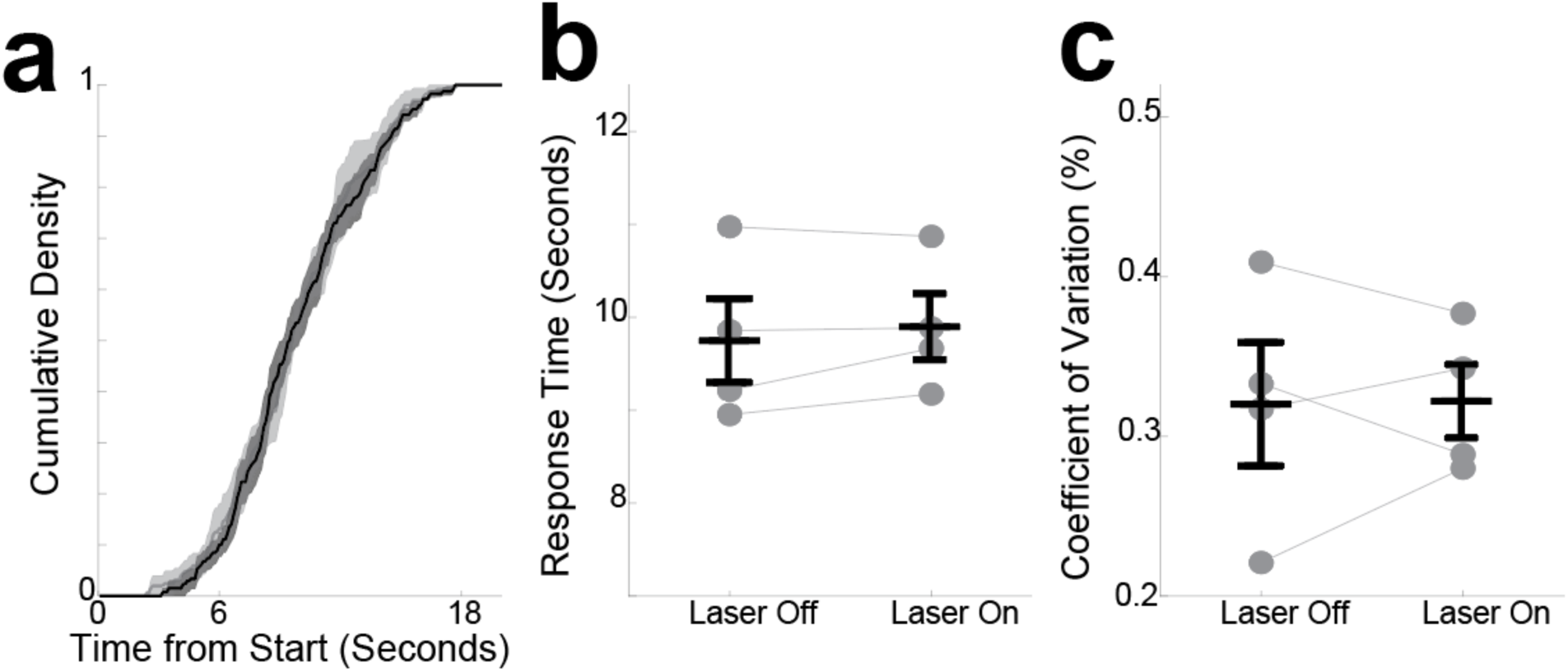
Optogenetic control mice. a) We expressed AAV-GFP in the PFC and implanted fiber optics in the DMS or the MD, as in Figure 4. On trials with the Laser On, we found a) no reliable effects on cumulative densities of response time, b) average response times, or c) coefficients of variation.

**Figure S4:**
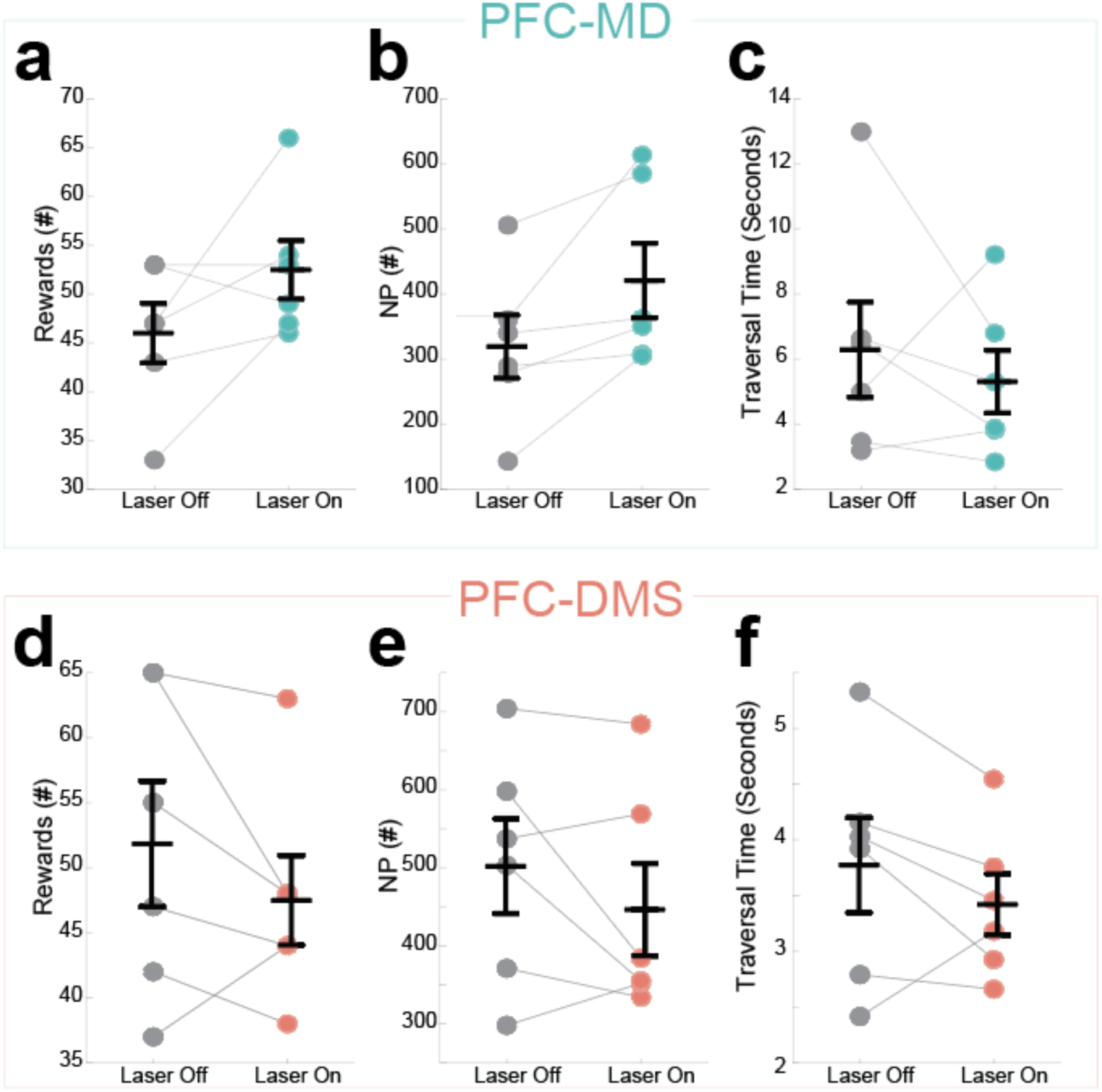
Optogenetic effects on rewards and response rates. For PFC-MD inactivation, we found no reliable difference in a) the number of rewards, b) the number of nosepokes, or c) the time between leaving the short nosepoke and arriving at the long nosepoke (traversal time), a measure of motivation of movement. We also found no differences for PFC-DMS inactivation in a)the number of rewards, b) the number of nosepokes, or c) traversal time. Same mice as in Figure 4.

**Figure S5:**
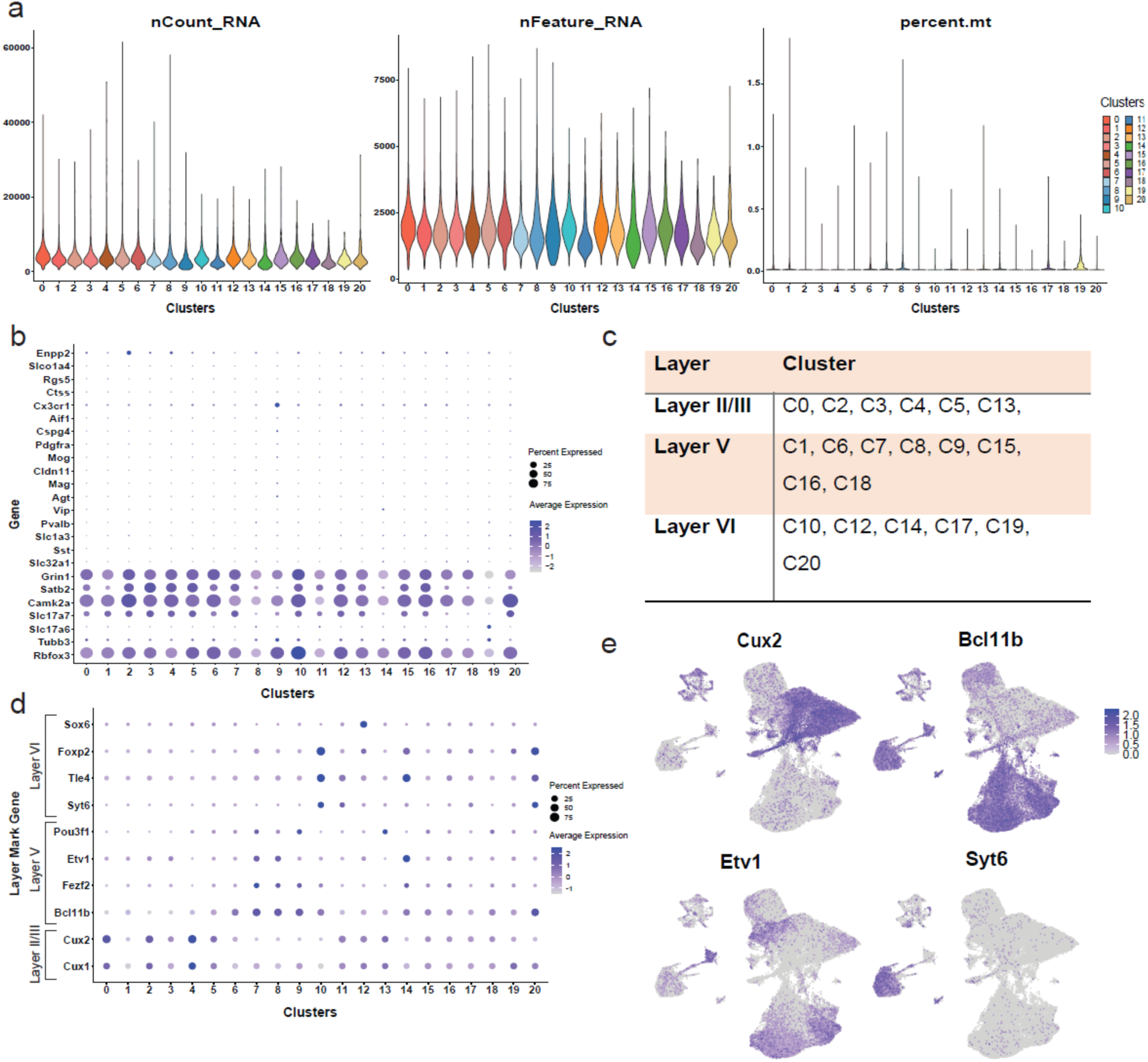
Quality control for snRNA-seq. a) Violin plots of scRNA-seq quality control metrics: the distribution of total RNA counts (nCount_RNA); detected genes (nFeature_RNA); and mitochondrial gene percentage (percent.mt) across cell clusters. b) Dotplot of neuronal marker genes. c) Table summarizing the layer assignment of each cluster. d) Dotplot of layer marker genes. e) Expression of specific markers of cortical layers.

**Figure S6:**
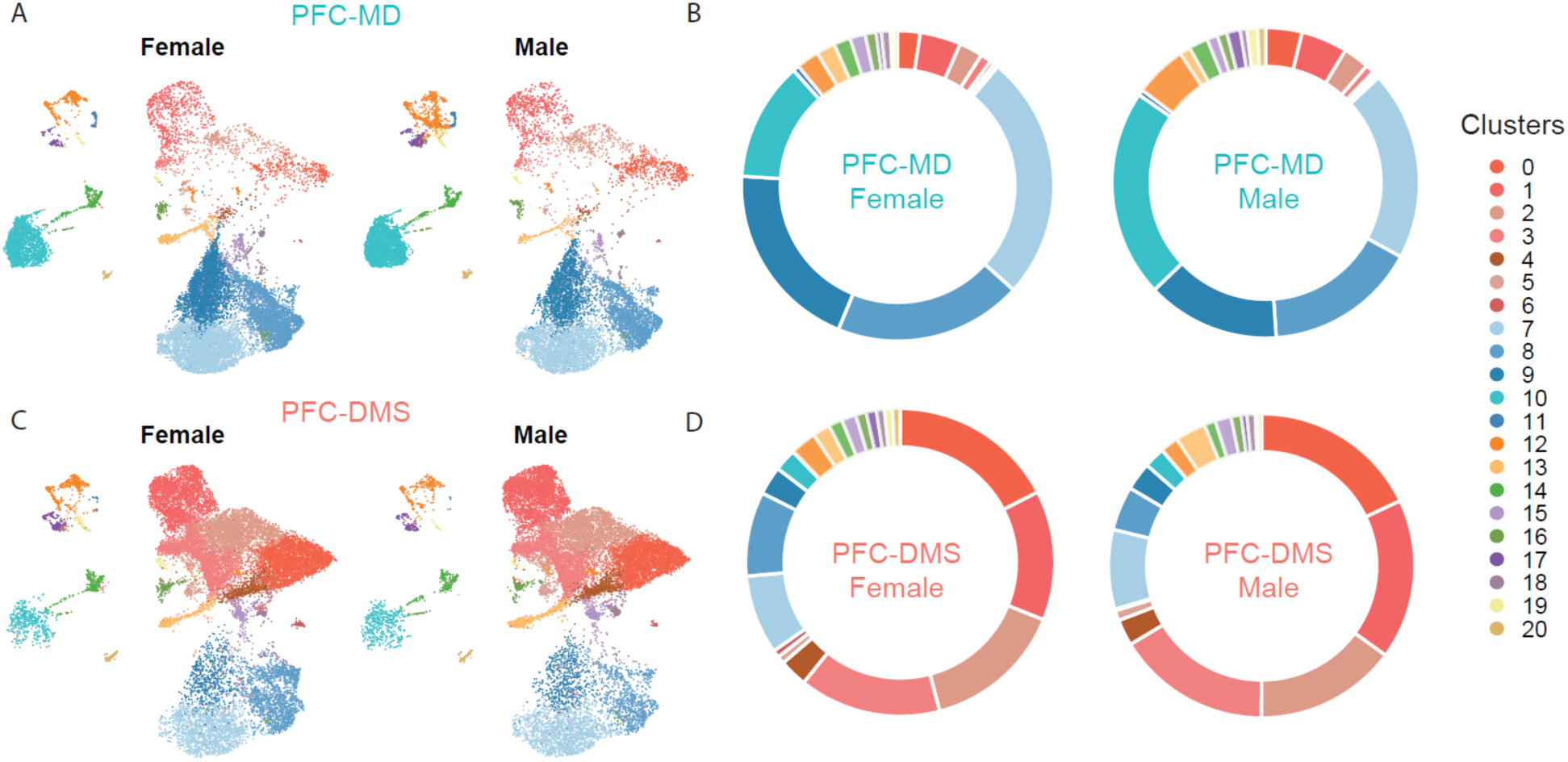
Sex effects. a) Uniform Manifold Approximation and Projection (UMAP) visualization of the expression of 21 clusters in PFC-DMS neurons from female and male mice. b) The percentage of each cluster in PFC-DMS neurons from female and male mice is similar, except that there is a higher percentage of clusters 6, 11, 19, and 20 in females compared to males. c) UMAP visualization of the expression of 21 clusters in PFC-MD neurons from female and male mice. d) The percentage of each cluster in PFC-MD neurons from female and male mice is similar, except that there is a higher percentage of clusters 12, 19, and 20 in males compared to females.

## References

1. Friedman, N. P. & Robbins, T. W. The role of prefrontal cortex in cognitive control and executive function. Neuropsychopharmacol. 47, 72–89 (2022).

2. Shallice, T. & Burgess, P. W. DEFICITS IN STRATEGY APPLICATION FOLLOWING FRONTAL LOBE DAMAGE IN MAN. Brain 114, 727–741 (1991).

3. Picton, T. W., Stuss, D. T., Shallice, T., Alexander, M. P. & Gillingham, S. Keeping time: effects of focal frontal lesions. Neuropsychologia 44, 1195–209 (2006).

4. Ma, S. et al. Molecular and cellular evolution of the primate dorsolateral prefrontal cortex. Science 377, eabo7257 (2022).

5. Narayanan, N. S. & Albin, R. L. Cognition in Parkinson’s Disease. (Elsevier, Acad. Press, 2022).

6. Goldman-Rakic, P. S., Castner, S. A., Svensson, T. H., Siever, L. J. & Williams, G. V. Targeting the dopamine D1 receptor in schizophrenia: insights for cognitive dysfunction. Psychopharmacology (Berl.) 174, 3–16 (2004).

7. Ährlund-Richter, S. et al. A whole-brain atlas of monosynaptic input targeting four different cell types in the medial prefrontal cortex of the mouse. Nat Neurosci 22, 657–668 (2019).

8. Gabbott, P. L. A., Warner, T. A., Jays, P. R. L., Salway, P. & Busby, S. J. Prefrontal cortex in the rat: projections to subcortical autonomic, motor, and limbic centers. J. Comp. Neurol 492, 145–177 (2005).

9. Anastasiades, P. G. & Carter, A. G. Circuit organization of the rodent medial prefrontal cortex. Trends in Neurosciences 44, 550–563 (2021).

10. Otis, J. M. et al. Prefrontal cortex output circuits guide reward seeking through divergent cue encoding. Nature 543, 103–107 (2017).

11. de Kloet, S. F. et al. Bi-directional regulation of cognitive control by distinct prefrontal cortical output neurons to thalamus and striatum. Nat Commun 12, 1994 (2021).

12. Wilhelm, M. et al. Striatum-projecting prefrontal cortex neurons support working memory maintenance. Nat Commun 14, 7016 (2023).

13. Buhusi, C. V. & Meck, W. H. What makes us tick? Functional and neural mechanisms of interval timing. Nat. Rev. Neurosci 6, 755–765 (2005).

14. Kim, J., Jung, A. H., Byun, J., Jo, S. & Jung, M. W. Inactivation of medial prefrontal cortex impairs time interval discrimination in rats. Front Behav Neurosci 3, 38 (2009).

15. Narayanan, N. S., Land, B. B., Solder, J. E., Deisseroth, K. & DiLeone, R. J. Prefrontal D1 dopamine signaling is required for temporal control. Proc. Natl. Acad. Sci. U.S.A. 109, 20726–20731 (2012).

16. Xu, M., Zhang, S., Dan, Y. & Poo, M. Representation of interval timing by temporally scalable firing patterns in rat prefrontal cortex. Proc. Natl. Acad. Sci. U.S.A. 111, 480–485 (2014).

17. Emmons, E. B. et al. Rodent Medial Frontal Control of Temporal Processing in the Dorsomedial Striatum. J. Neurosci. 37, 8718–8733 (2017).

18. Wang, J., Narain, D., Hosseini, E. A. & Jazayeri, M. Flexible timing by temporal scaling of cortical responses. Nature Neuroscience 21, 102 (2018).

19. Singh, A. et al. Timing variability and midfrontal ∼4 Hz rhythms correlate with cognition in Parkinson’s disease. NPJ Parkinsons Dis 7, 14 (2021).

20. Parker, K. L. et al. Delta-frequency stimulation of cerebellar projections can compensate for schizophrenia-related medial frontal dysfunction. Mol Psychiatry 22, 647–655 (2017).

21. Ward, R. D., Kellendonk, C., Kandel, E. R. & Balsam, P. D. Timing as a window on cognition in schizophrenia. Neuropharmacology (2011) doi:10.1016/j.neuropharm.2011.04.014.

22. Merchant, H. & de Lafuente, V. Introduction to the neurobiology of interval timing. Adv. Exp. Med. Biol. 829, 1–13 (2014).

23. Narayanan, N. S. Ramping activity is a cortical mechanism of temporal control of action. Curr Opin Behav Sci 8, 226–230 (2016).

24. Bakhurin, K. I. et al. Differential Encoding of Time by Prefrontal and Striatal Network Dynamics. J. Neurosci. 37, 854–870 (2017).

25. Treisman, M. Temporal discrimination and the indifference interval: Implications for a model of the ‘internal clock’. Psychological Monographs: General and Applied 77, 1–31 (1963).

26. Simen, P., Balci, F., de Souza, L., Cohen, J. D. & Holmes, P. A model of interval timing by neural integration. J. Neurosci. 31, 9238–9253 (2011).

27. Bruce, R. A. et al. Complementary cognitive roles for D2-MSNs and D1-MSNs during interval timing. eLife 13, RP96287 (2025).

28. Kononowicz, T. W. & van Rijn, H. Decoupling interval timing and climbing neural activity: a dissociation between CNV and N1P2 amplitudes. J Neurosci 34, 2931–2939 (2014).

29. Kononowicz, T. W. & Van Rijn, H. Slow Potentials in Time Estimation: The Role of Temporal Accumulation and Habituation. Front. Integr. Neurosci. 5, (2011).

30. Kim, Y.-C. et al. Optogenetic Stimulation of Frontal D1 Neurons Compensates for Impaired Temporal Control of Action in Dopamine-Depleted Mice. Curr Biol 27, 39–47 (2017).

31. Stutt, H. R. et al. Sex similarities and dopaminergic differences in interval timing. Behav Neurosci 138, 85–93 (2024).

32. Balci, F. et al. Interval timing in genetically modified mice: a simple paradigm. Genes Brain Behav 7, 373–384 (2008).

33. Tosun, T., Gur, E. & Balci, F. Mice plan decision strategies based on previously learned time intervals, locations, and probabilities. Proc Natl Acad Sci U S A 113, 787–792 (2016).

34. Bruce, R. A. et al. Experience-related enhancements in striatal temporal encoding. Eur J Neurosci 54, 5063–5074 (2021).

35. Bruce, R. A. et al. Complementary cognitive roles for D2-MSNs and D1-MSNs in interval timing. bioRxiv 2023.07.25.550569 (2023) doi:10.1101/2023.07.25.550569.

36. Larson, T. et al. Mice expressing P301S mutant human tau have deficits in interval timing. Behav Brain Res 432, 113967 (2022).

37. Weber, M. A. et al. Glycolysis-enhancing α1-adrenergic antagonists modify cognitive symptoms related to Parkinson’s disease. npj Parkinsons Dis. 9, 1–7 (2023).

38. Han, S.-W., Kim, Y.-C. & Narayanan, N. S. Projection targets of medial frontal D1DR-expressing neurons. Neuroscience Letters 655, 166–171 (2017).

39. Lusk, N. A., Petter, E. A. & Meck, W. H. A systematic exploration of temporal bisection models across sub- and supra-second duration ranges. Journal of Mathematical Psychology 94, 102311 (2020).

40. Merchant, H. et al. Sensorimotor neural dynamics during isochronous tapping in the medial premotor cortex of the macaque. Eur J Neurosci 41, 586–602 (2015).

41. Chapin, J. K. & Nicolelis, M. A. Principal component analysis of neuronal ensemble activity reveals multidimensional somatosensory representations. J. Neurosci. Methods 94, 121–140 (1999).

42. Narayanan, N. S. & Laubach, M. Delay activity in rodent frontal cortex during a simple reaction time task. J. Neurophysiol 101, 2859–2871 (2009).

43. Parker, K. L., Chen, K.-H., Kingyon, J. R., Cavanagh, J. F. & Narayanan, N. S. D1-Dependent 4 Hz Oscillations and Ramping Activity in Rodent Medial Frontal Cortex during Interval Timing. J. Neurosci. 34, 16774–16783 (2014).

44. Emmons, E. B. et al. Temporal Learning Among Prefrontal and Striatal Ensembles. Cerebral Cortex Communications 1, (2020).

45. Miškić, T., Kostović, I., Rašin, M.-R. & Krsnik, Ž. Adult Upper Cortical Layer Specific Transcription Factor CUX2 Is Expressed in Transient Subplate and Marginal Zone Neurons of the Developing Human Brain. Cells 10, 415 (2021).

46. Franco, S. J. et al. Fate-Restricted Neural Progenitors in the Mammalian Cerebral Cortex. Science 337, 746–749 (2012).

47. Arlotta, P. et al. Neuronal Subtype-Specific Genes that Control Corticospinal Motor Neuron Development In Vivo. Neuron 45, 207–221 (2005).

48. Du, H. et al. Transcription factors Bcl11a and Bcl11b are required for the production and differentiation of cortical projection neurons. Cereb Cortex 32, 3611–3632 (2021).

49. Guan, D., Armstrong, W. E. & Foehring, R. C. Electrophysiological properties of genetically identified subtypes of layer 5 neocortical pyramidal neurons: Ca2+ dependence and differential modulation by norepinephrine. J Neurophysiol 113, 2014–2032 (2015).

50. Vaasjo, L. O. et al. Characterization and manipulation of Corticothalamic neurons in associative cortices using Syt6-Cre transgenic mice. Journal of Comparative Neurology 530, 1020–1048 (2022).

51. Molyneaux, B. J., Arlotta, P., Menezes, J. R. L. & Macklis, J. D. Neuronal subtype specification in the cerebral cortex. Nat Rev Neurosci 8, 427–437 (2007).

52. Arnsten, A. F. T. Catecholamine influences on dorsolateral prefrontal cortical networks. Biol. Psychiatry 69, e89–99 (2011).

53. Narayanan, N. S., Rodnitzky, R. L. & Uc, E. Y. Prefrontal dopamine signaling and cognitive symptoms of Parkinson’s disease. Rev Neurosci 24, 267–278 (2013).

54. Singh, A. et al. Gene regulatory landscape of cerebral cortex folding. Sci Adv 10, eadn1640 (2024).

55. Barington, M., Risom, L., Ek, J., Uldall, P. & Ostergaard, E. A recurrent de novo CUX2 missense variant associated with intellectual disability, seizures, and autism spectrum disorder. Eur J Hum Genet 26, 1388–1391 (2018).

56. Astudillo, D. et al. CaMKII inhibitor 1 (CaMK2N1) mRNA is upregulated following LTP induction in hippocampal slices. Synapse 74, e22158 (2020).

57. Lieberman, P. FOXP2 and Human Cognition. Cell 137, 800–802 (2009).

58. Lin, L.-C., Cole, R. C., Greenlee, J. D. W. & Narayanan, N. S. A Pilot Study of Ex Vivo Human Prefrontal RNA Transcriptomics in Parkinson’s Disease. Cell Mol Neurobiol 43, 3037–3046 (2023).

59. Miller, E. K. The Prefrontal Cortex: Complex Neural Properties for Complex Behavior. Neuron 22, 15–17 (1999).

60. Stine, G. M. & Jazayeri, M. Control Principles of Neural Dynamics Revealed by the Neurobiology of Timing. Annu Rev Neurosci (2025) doi:10.1146/annurev-neuro-091724-015512.

61. Emmons, E. B., Kennedy, M., Kim, Y. & Narayanan, N. S. Corticostriatal stimulation compensates for medial frontal inactivation during interval timing. Sci Rep 9, 14371 (2019).

62. Lusk, N., Meck, W. H. & Yin, H. H. Mediodorsal Thalamus Contributes to the Timing of Instrumental Actions. J Neurosci 40, 6379–6388 (2020).

63. Hintiryan, H. et al. The mouse cortico-striatal projectome. Nat Neurosci 19, 1100–1114 (2016).

64. Yao, Z. et al. A taxonomy of transcriptomic cell types across the isocortex and hippocampal formation. Cell 184, 3222–3241.e26 (2021).

65. Bhattacherjee, A. et al. Cell type-specific transcriptional programs in mouse prefrontal cortex during adolescence and addiction. Nat Commun 10, 4169 (2019).

66. Bhattacherjee, A. et al. Spatial transcriptomics reveals the distinct organization of mouse prefrontal cortex and neuronal subtypes regulating chronic pain. Nat Neurosci 26, 1880–1893 (2023).

67. Yao, Z. et al. A high-resolution transcriptomic and spatial atlas of cell types in the whole mouse brain. Nature 624, 317–332 (2023).

68. Gao, L. et al. Single-neuron projectome of mouse prefrontal cortex. Nat Neurosci 25, 515–529 (2022).

69. Murphy, S. E., Wright, L. C., Browning, M., Cowen, P. J. & Harmer, C. J. A role for 5-HT4 receptors in human learning and memory. Psychological Medicine 50, 2722–2730 (2020).

70. Hisaoka, T., Nakamura, Y., Senba, E. & Morikawa, Y. The forkhead transcription factors, Foxp1 and Foxp2, identify different subpopulations of projection neurons in the mouse cerebral cortex. Neuroscience 166, 551–563 (2010).

71. Kleiner, M., Brainard, D. & Pelli, D. What’s new in Psychtoolbox-3? (2007).

72. Singh, A. et al. Evoked mid-frontal activity predicts cognitive dysfunction in Parkinson’s disease. J Neurol Neurosurg Psychiatry jnnp-2022-330154 (2023) doi:10.1136/jnnp-2022-330154.

73. Cole, R. C. et al. Novelty-induced frontal-STN networks in Parkinson’s disease. Cereb Cortex 33, 469–485 (2022).

74. Roh, H. C. et al. Simultaneous Transcriptional and Epigenomic Profiling from Specific Cell Types within Heterogeneous Tissues In Vivo. Cell Rep 18, 1048–1061 (2017).

75. Weber, M. A. et al. Amphetamine increases timing variability by degrading prefrontal ramping activity. 2024.09.26.615252 Preprint at 10.1101/2024.09.26.615252 (2024).

76. Pan, C. et al. Shrinkage-mediated imaging of entire organs and organisms using uDISCO. Nat Methods 13, 859–867 (2016).

77. Benjamini, Y. & Hochberg, Y. Controlling the false discovery rate: a practical and powerful approach to multiple testing. Journal of the Royal Statistical Society. Series B (Methodological) 289–300 (1995).

78. Simpson, E. H. et al. Lights, fiber, action! A primer on in vivo fiber photometry. Neuron 112, 718–739 (2024).

79. Kim YC, Miller A, Lins LC, Han SW, Keiser MS, Boudreau RL, Davidson BL, Narayanan NS. RNA Interference of Human α-Synuclein in Mouse. Front Neurol. 2017 Jan 31;8:13. doi: 10.3389/fneur.2017.00013. PMID: 28197125; PMCID: PMC5281542.

80. Schwalbe, D. C. et al. Molecular Organization of Autonomic, Respiratory, and Spinally-Projecting Neurons in the Mouse Ventrolateral Medulla. J Neurosci 44, e2211232024 (2024).

81. Fleming, S. J. et al. Unsupervised removal of systematic background noise from droplet-based single-cell experiments using CellBender. Nature Methods 20, 1323–1335 (2023).

82. Hao, Y. et al. Dictionary learning for integrative, multimodal and scalable single-cell analysis. Nature Biotechnology 42, 293–304 (2024).

83. Tsyporin, J. et al. Transcriptional repression by FEZF2 restricts alternative identities of cortical projection neurons. Cell Rep 35, 109269 (2021).

84. Lambe, E. K., Fillman, S. G., Webster, M. J. & Shannon Weickert, C. Serotonin receptor expression in human prefrontal cortex: balancing excitation and inhibition across postnatal development. PLoS One 6, e22799 (2011).

